# ADT-1004: A First-in-Class, Orally Bioavailable Selective pan-RAS Inhibitor for Pancreatic Ductal Adenocarcinoma

**DOI:** 10.1101/2024.10.04.616725

**Authors:** Dhana Sekhar Reddy Bandi, Ganji Purnachandra Nagaraju, Sujith Sarvesh, Julienne L. Carstens, Jeremy B. Foote, Emily C. Graff, Yu-Hua D Fang, Adam B. Keeton, Xi Chen, Kristy L. Berry, Sejong Bae, Mehmet Akce, Greg Gorman, Karina J. Yoon, Upender Manne, Micheal R. Boyd, Donald J. Buchsbaum, Asfar S. Azmi, Yulia Y. Maxuitenko, Gary A. Piazza, Bassel F. El-Rayes

**Author notes:** **Corresponding Author:** Bassel F. El-Rayes, MD Albert F. LoBuglio Endowed Chair for Translational Cancer Research, Division Director, Hematology and Oncology, Deputy Director, O’Neal Comprehensive Cancer Center, Heersink School of Medicine, University of Alabama at Birmingham. **Disclosure of Potential Conflicts of Interest:** G.A. Piazza, A.B. Keeton, X. Chen, and M.R. Boyd are co-founders of ADT Pharmaceuticals Inc. and consultants. All other authors declare no potential conflicts of interest. **Funding Sources:** NIH/NCI R01CA254197, R01CA238514, and P30CA01314.

## Abstract

Here, we evaluated *in vivo* antitumor activity, target engagement, selectivity, and tumor specificity of ADT-1004, an orally bioavailable prodrug of ADT-007 having highly potent and selective pan-RAS inhibitory activity. ADT-1004 strongly blocked tumor growth and RAS activation in mouse PDAC models without discernable toxicity. As evidence of target engagement and tumor specificity, ADT-1004 inhibited activated RAS and ERK phosphorylation in PDAC tumors at dosages approximately 10-fold below the maximum tolerated dose and without discernable toxicity. ADT-1004 inhibited ERK phosphorylation in PDAC tumors. In addition, ADT-1004 blocked tumor growth and ERK phosphorylation in PDX PDAC models with KRAS^G12D^, KRAS^G12V^, KRAS^G12C^, or KRAS^G13Q^ mutations. ADT-1004 treatment increased CD4^+^ and CD8^+^ T cells in the TME consistent with exhaustion and increased MHCII^+^ M1 macrophage and dendritic cells. ADT-1004 demonstrated superior efficacy over sotorasib and adagrasib in tumor models involving human PDAC cells resistant to these KRAS^G12C^ inhibitors. As evidence of selectivity for tumors from PDAC cells with mutant KRAS, ADT-1004 did not impact the growth of tumors from RAS^WT^ PDAC cells. Displaying broad antitumor activity in multiple mouse models of PDAC, along with target engagement and selectivity at dosages that were well tolerated, ADT-1004 warrants further development.

**Significance:** ADT-1004 displayed robust antitumor activity in aggressive and clinically relevant PDAC models with unique tumor specificity to block RAS activation and MAPK signaling in RAS mutant cells. As a pan-RAS inhibitor, ADT-1004 has broad activity and potential efficacy advantages over allele-specific KRAS inhibitors by averting resistance. These findings support clinical trials of ADT-1004 for KRAS mutant PDAC.

## Introduction

Pancreatic ductal adenocarcinoma (PDAC) is projected to become the second leading cause of cancer deaths in the US by 2040.^1, 2^ In 2024, PDAC is expected to cause 52,580 deaths in the US.^3^ Systemic treatment strategies for PDAC have yielded little improvement in survival.^4^ The overall survival rate at 5-years remains at 12% for patients diagnosed at early stages or 3% for those diagnosed with metastatic disease.^5^ Hence, developing novel systemic treatment strategies is essential for improving the survival of patients with PDAC.

The gene encoding for KRAS, a small GTPase protein, is mutated in more than 90% of PDAC patients. KRAS mutations in patients with PDAC include G12D (37.0%), G12V (28.2%), G12R (12.7%), G12C (2.7%), or others (7.0%), and some have multiple mutations in RAS (2.1%).^6,7,8^ The National Cancer Institute has identified anti-KRAS testing as one of four essential components for PDAC drug screening and development.^9,10^ Targeting this oncoprotein has proven challenging because it lacks a relatively flat surface for suitable small molecule binding sites.^11^ To date, the U.S. Food and Drug Administration (FDA) has approved two KRAS^G12C^ inhibitors, sotorasib (AMG-510) and adagrasib (MRTX849) for treatment of lung cancer. Several newer inhibitors targeting different KRAS mutations are in preclinical development or clinical trials. Sotorasib and adagrasib have limited use in PDAC, given that KRAS^G12C^ is mutated in less than 3% of patients diagnosed with PDAC.^12^ The development of adaptive resistance also limits the efficacy KRAS^G12C^ inhibitors in patients diagnosed with KRAS^G12C^ mutant cancers.^13^ Resistance mechanisms include new KRAS mutations or activation of RAS^WT^ isozymes (HRAS and NRAS) from upstream receptor tyrosine kinases.^14^ Consequently, a pan-RAS inhibitor could provide a more effective approach to address the complex mutational landscape of PDAC that can escape compensatory mechanisms contributing to intrinsic or adaptive resistance.

To address these limitations, we developed and recently characterized ADT-007, a highly potent and selective pan-RAS inhibitor with broad activity against a histologically diverse range of RAS-mutant cancer cell lines.^15^ ADT-1004 is an orally bioavailable prodrug of ADT-007 designed to improve the water solubility and metabolic stability of ADT-007 (**Figure 1A**). This study evaluated the antitumor activity and target selectivity of ADT-1004 in aggressive and clinically relevant mouse and PDX models of PDAC. We also defined various pharmacological attributes of ADT-1004 that could set this drug candidate apart from other KRAS inhibitors approved or in development.

**Figure 1:**
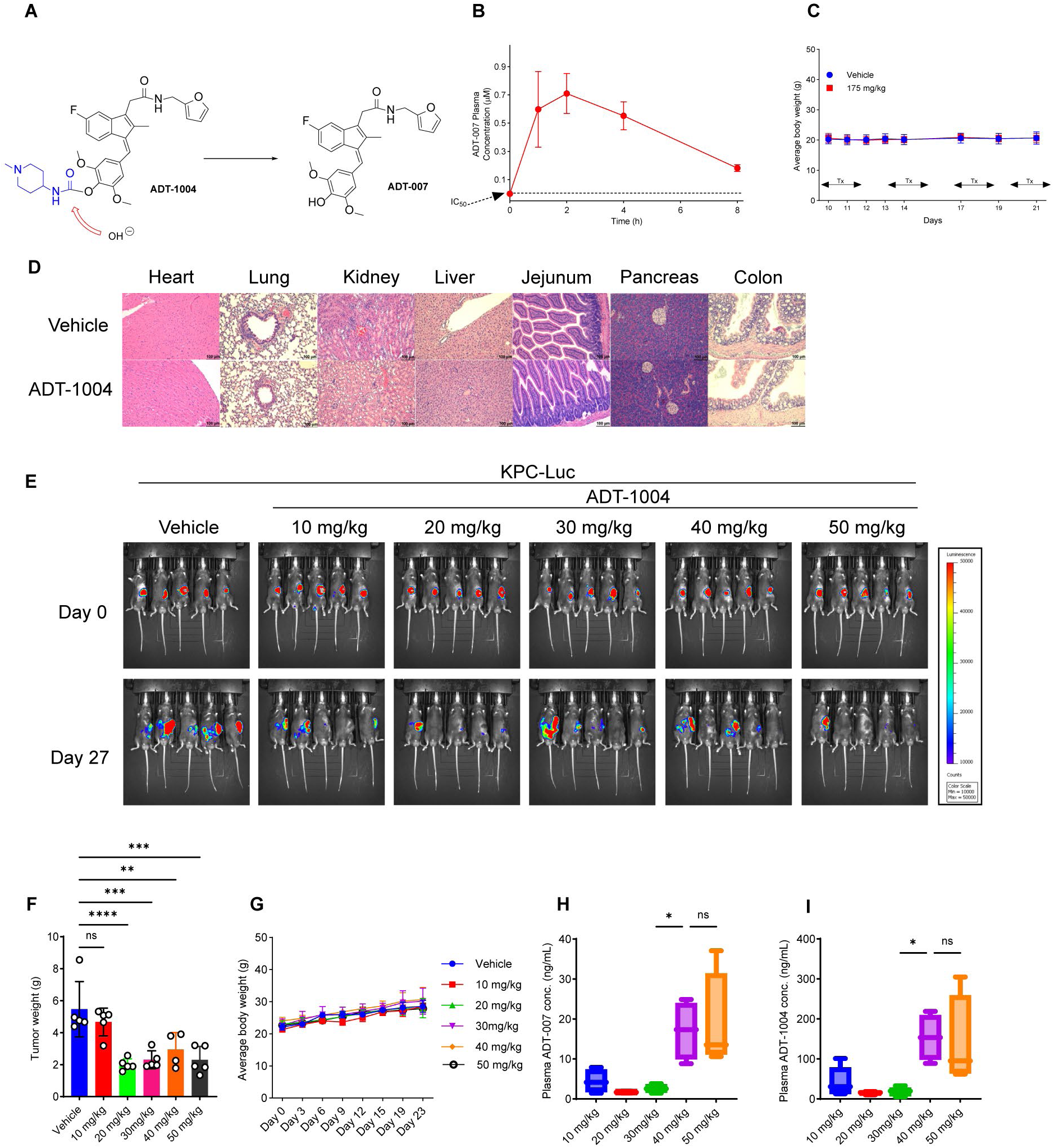
Evaluation of ADT-1004 efficacy and toxicity profile in murine model of pancreatic cancer. (**A**). Chemical structures of ADT-1004 and ADT-007. **(B).** Plasma levels of ADT-007 in mice following a single oral administration of ADT-1004 (100 mg/kg, mean ± SD, n=3-4). **(C).** Body weights of C57BL/6 mice treated with ADT-1004 (175 mg/kg, n=5) or vehicle (n=10) orally BID, 5days/week for 3.5 weeks (mean ± SD). (**D**). Histopathological examination of vital organs (heart, lung, kidney, liver, jejunum, pancreas, and colon) from mice treated with ADT-1004 at 175 mg/kg BID as assessed by H&E staining. (**E**). *F*-Luc-labeled KPC cells were orthotopically implanted into C57BL/6J mice (n=30). After 1 week, group of 5 mice were orally administered vehicle or ADT-1004 (10, 20, 30, 40, and 50 mg/kg). Bioluminescence images at the indicated time points are shown. (**F**). Tumors were weighed at the end of the experiment under the indicated conditions for experiment in (**E**). (**G**). Average body weights of the animals during the course of the treatment shown in experiment (**E**). (**H**). The concentration of ADT-007 in plasma of mice on the last day of treatment is depicted as mean alongside SD. (**I**). The concentration of ADT-1004 in plasma of mice on the last day of treatment is depicted as geometric mean alongside standard deviation. ADT-007 and ADT-1004 levels measured by LC-MS/MS. All quantitative data represent the mean ± SEM. ns, non-significant, ∗p < 0.05, ∗∗p < 0.01, ∗∗∗p < 0.001, and ∗∗∗∗p < 0.0001.

## Materials and methods

### Synthesis of ADT-1004

Synthesis of ADT-1004 HCl [(Z)-4-((5-fluoro-3-(2-((furan-2-ylmethyl)amino)-2-oxoethyl)-2-methyl-1H-inden-1-ylidene)methyl)-2,6-dimethoxyphenyl (1-methylpiperidin-4-yl)carbamate hydrogen chloride]: To a solution of 1-methylpiperidin-4-amine (2.3 g, 20 mmol) in dry chloroform (40 mL, freshly distilled from P_2_O_5_), triphosgene (2.2 g, 7 mmol) was added in portion with stirring, followed by addition of triethylamine (7 mL). The reaction mixture was refluxed for 6h. (Z)-2-(5-fluoro-1-(4-hydroxy-3,5-dimethoxybenzylidene)-2-methyl-1H-inden-3-yl)-N-(furan-2-ylmethyl) acetamide (ADT-007, synthesis was previously described ^15^) (3.0 g, 6.7 mmol) and dibutyltin dilaurate (7.4 g, 11.6 mmol) were added and the mixture synthesized from the reaction was stirred at 65 °C overnight. The solvents were extracted under vacuum and silica gel column chromatography was used to purify the residue, followed by the addition of 1.00 eq. of HCl (1N in dioxane, 0 °C) to yield ADT-1004 HCl as a yellow crystal (yield was 55%).

### Cell lines

The human PDAC cell lines, MIA PaCa-2 and BxPc-3, were acquired from the American Type Culture Collection (ATCC; Manassas, VA, USA) and cultured under the manufacturer’s provided guidelines. 2838c3 mouse PDAC cells were purchased from Kerafast (Boston, MA, USA), and the KPC cell line was gifted by Dr. Gregory Lesinski (Emory University, USA). MIA PaCa-2 cells resistant to KRAS^G12C^-inhibitors (MIA-AMG-Res) were generated by us as previously described.^16^ All cells were grown in a humidified atmosphere at 37 °C with 5% CO_2_, utilizing either Dulbecco’s Modified Eagle Medium (DMEM, ATCC, #30-2002) for 2838c3 or Roswell Park Memorial Institute (RPMI)-1640 Medium (ATCC, #30-2001) for BxPc3 with 10% fetal bovine serum (FBS, ATCC, #30-2020) and 1% penicillin/streptomycin (ATCC, #30-2101) in common. Furthermore, MIA PaCa-2 cells received an additional supplementation of 1% horse serum (Gibco, #16050-122) in DMEM along with 10% FBS and 1% penicillin/streptomycin. All cell lines were routinely tested for mycoplasma and other pathogen contaminations.

### Cell growth assays

MIA PaCa-2 and MIA-AMG-Res were plated in 96-well culture plates at a density of 4×10^3^ cells per well. After overnight incubation, the cells were treated with ADT-007, sotorasib, or adagrasib (Selleckchem, Houston, TX, USA). After 72 h, 10 μl methylthiazole tetrazolium (MTT; 5 mg/mL in PBS; Sigma-Aldrich) was added, and the cells were incubated for another 2 h at 37 °C. The resulting formazan crystals were solubilized in 100 μl DMSO and mixed well by pipetting, and absorbance was measured at 570 nm and 630 nm using the Biotek Synergy MX Multi Format Microplate Reader. The average measurement at 630 nm was subtracted from the average at 590 nm, and relative growth rate was plotted with respect to control-treated cells.

### Clonogenic assays

At a density of 1000 cells per well in 6-well plates, MIA PaCa-2 and MIA-AMG-Res cells were seeded and exposed to ADT-007, sotorasib, or adagrasib for 72 h. Following the treatment, the medium containing the drug was replaced with fresh medium, and the plates were then placed in a CO_2_ incubator for 1 week (for MIA-AMG-Res colonies) or 10 days (for MIA PaCa-2 colonies). After the incubation period, the medium was removed, and colonies were fixed using methanol, followed by crystal violet staining for 15 min. The colonies were photographed after a thorough wash and drying process.

### Immunoblot analysis

Tumor tissues were lysed in RIPA buffer (Thermo Scientific, #89901) and protein concentrations assayed using a bicinchoninic acid (BCA) protein assay (Thermo Fisher Scientific, #23225). 40 μg of protein lysate was separated through 4-20% SDS-PAGE (BIO-RAD, #4568096) and transferred onto PVDF membranes (Invitrogen, #IB34001). The membranes were then incubated with antibodies diluted in 2% Bovine Serum Albumin (BSA, Fisher Scientific, #BP1600) for 2 h at room temperature. Primary antibodies were phospho ERK (Cell Signaling Technology #9121; 1:1000), total ERK (Cell Signaling Technology (CST) #9122; 1:1000), anti-beta actin (Santa Cruz #sc-47724; 1:3,000). Incubation with HRP-linked secondary antibodies (CST, #7074/7076;) at a dilution of 1:3000 in a 2% BSA solution was carried out for 1 h at room temperature. The signal was then detected on the LI-COR Odyssey DLx Imager using the ECL chemiluminescence detection system (Thermo Fisher Scientific, #34577).

### Active RAS detection assay

RAS activation (RAS-GTP) levels were measured using the Cell Signaling Technology RAS activation assay kit (#8821). Lysates were prepared from tumors collected from ADT-1004 or vehicle treated mice by performing the steps provided by the manufacturer protocol. Briefly, tumors were lysed using the provided lysis buffer supplemented with protease and phosphatase inhibitors. A total of 1 mg/mL lysate in 1X lysis buffer was employed for the experiment. Equal amounts of protein were then incubated with the GST-Raf1-RBD protein, and the reaction mixture was loaded onto a RAS affinity resin to capture activated RAS. Following extensive washing to remove unbound proteins, the activated RAS was eluted in sample buffer and subjected to immunoblotting using Ras Mouse mAb (1:200 dilution) with gentle agitation overnight at 4 °C. The membrane was then probed with anti-mouse IgG, HRP-linked antibody (1:2000, #7076), and HRP-conjugated anti-biotin antibody (1:1000, #7075) to detect biotinylated protein markers. The signal was detected using chemiluminescent reagents, and the resulting bands were quantified using an imaging system.

### Pharmacokinetic study

Following acclimatization in the Biologic Research Laboratory at the University of South Alabama, College of Medicine 6-8-week-old female C57BL/6 mice (Envigo, strain 044) were randomly assigned to two groups (n=4) and received ADT-1004 (100 mg/kg, in water) by oral gavage once. Blood was drawn at 1, 2, 4, or 8 h (each animal was bled twice) into K_2_EDTA-microtainer tubes followed by separation of plasma. ADT-1004 levels in plasma as well as ADT-007 levels generated following oral administration of ADT-1004 were determined using reverse phase chromatography with MS/MS detection. The study followed established guidelines and adhered to the approved protocol of the University of South Alabama Institutional Animal Care and Use Committee (IACUC).

### Maximum tolerated dose (MTD) study

Following acclimatization in Biological Research Facility at Auburn University 7-8-week-old female C57BL/6 mice (Charles River Laboratories, stain 027) which were subcutaneously implanted with mouse 2838c3 PDAC cells (1 × 10^6^ cells resuspended in 100 μL PBS), were randomly assigned to groups and received either vehicle (citric acid with sodium citrate monobasic in water, n=10) or ADT-1004 (125-225 mg/kg, n=5/dose) by oral gavage twice daily (BID), five days a week (5x/week) for 3.5 weeks. Mice were observed for adverse reaction to the treatment daily and tumor measurements and body weights were monitored biweekly. At the end of the study (a total of 18 daily treatments), mice in the group treated with MTD of ADT-1004 (defined as a dose that produced no deaths and no more than 5% body weight loss) were euthanized, and blood was collected and submitted for clinical chemistries and complete blood counts to the Auburn University Clinical Pathology Laboratory. Mice were necropsied, organs were collected and fixed for histopathological analysis, and bone marrow smears were prepared for cytology. Blinded assessment of organ viscera (heart, lung, kidney, liver, stomach, pancreas, duodenum, jejunum, ileum, cecum, and colon) was performed following standard procedures by board-certified veterinary pathologists. The MTD study followed established guidelines and adhered to the approved protocol of the Auburn University IACUC.

### Mouse tumor xenograft experiments

#### PDAC (MIA PaCa-2, MIA-AMG-Res, and BxPc-3) cell-derived tumor xenograft models

Following acclimatization in the University of Alabama at Birminghamn (UAB) animal housing facility, male NSG (NOD.Cg-Prkdc^scid^ Il2^rgtm1Wjl^/SzJ) mice aged 4 to 5 weeks (stock no. 005557; the Jackson Laboratory) were subcutaneously implanted with MIA PaCa-2, *MIA-AMG-Res*, or BxPc3 PDAC cells. A total of 1 × 10^6^ cells (for MIA PaCa-2 and *MIA-AMG-Res*) and 5 × 10^6^ cells (BxPc-3), resuspended in 200 μL PBS, were unilaterally injected into the right flank of mice using a BD 26Gx 5/8 1 mL Sub-Q syringe. Seven days post-transplantation or when tumors reached approximately 80–100 mm^3^ in size, recipient mice were randomly assigned to 4 groups (n=5) and received either vehicle, ADT-1004 (40 mg/kg), sotorasib (40 mg/kg), or adagrasib (40 mg/kg) through oral gavage. For BxPc3-implanted mice, two groups were established (n=5), and mice were orally administered either vehicle or ADT-1004 (40 mg/kg). Tumor volumes and body weights were monitored biweekly, with the formula to calculate the tumor volume as length × width^2^ × 0.5. After completing the drug dosing regimen, tumors from vehicle and treatment groups were utilized for molecular analysis. *In vivo* studies followed established guidelines and adhered to the approved protocol of the UAB Institutional Animal Care and Use Committee (IACUC).

#### Orthotopic pancreatic injection of murine PDAC cells (KPC and 2838c3)

C57BL6/J mice (stock no. 000664; The Jackson Laboratory) were subjected to isoflurane anesthesia, followed by an intra-abdominal incision to access the spleen and pancreas. A Matrigel suspension (40 μL), containing either KPC-*f*-luc or 2838c3-*f* -luc cells (1 × 10^5^), was injected into the tail of the pancreas. The skin and abdominal wall were then closed by suturing. Following a week, mice were divided into treatment groups at random (n=5-7 per group) and given oral dosages once daily (QD), 5 days/week of vehicle (citric acid with sodium citrate monobasic) or ADT-1004 (40 mg/kg) until completing the experiment. Imaging sessions involving D-luciferin injection were conducted bi-weekly throughout the study. The total luminescence from tumor-bearing regions was quantified using the Living image *in vivo* imaging software. No indications of toxicity, such as changes in body weight or abnormalities in organs, were observed. At the endpoint, mice were euthanized, and their tumors were harvested and weighed for analysis.

#### Flow cytometry

Tumors derived from 2838c3-*f-*luc and KPC-*f-*luc PDAC cells implanted into the pancreas of C57BL6/J mice were subjected to digestion using a solution comprising 0.1 mg/ml DNase 1 and 1 mg/ml collagenase IV (Worthington Biochemical, Lakewood, NJ) in Hank’s Balanced Salt Solution (HBSS) at 37 °C with shaking, for 45 min. Following digestion, the samples underwent rinsing with RPMI-1640 supplemented with 10% FBS, and a 70 μm strainer was used to filter them to produce single-cell suspensions. The separated cells were labeled for 30 to 60 min at 4°C using primary antibodies coupled to fluorophores and a live/dead dye (**Supplementary Table 1**). Cells were then rinsed and suspended in a flow buffer. After labeling the cell surface, the cells were fixed at 4 °C in 4% paraformaldehyde or FoxP3 transcription buffer set (eBioscience; # 00-5523-00) for 45 min and then washed with 1x Perm/Wash (BD, 554723). Data acquisition was conducted using a Symphony A5 flow cytometer, and analysis was performed using FlowJo version 10.7.2.

### KRAS –mutant patient-derived tumor xenograft studies

#### Mouse tumorigenesis experiments using PDAC PDX models

Mouse experiments were conducted using PDAC PDX models with distinct KRAS mutations G12D and G12C (generously provided by Dr. Karina Yoon), ^17^ G12V and G13Q (obtained from Dr. Upendar Manne). Male NSG mice (5–6 weeks old, stock no. 005557; the Jackson Laboratory) were utilized for the experiments. In brief, F1 generation tumors were cut into 2-mm × 2-mm fragments and subcutaneously implanted by way of a small incision produced in the right flanks of NSG mice while they were anesthetized. Tumor size and body weights were monitored biweekly. Tumor volume was calculated using the formula: length × width^^2^ × 0.5. Once tumors reached approximately 80–100 mm^^3^, mice (n=6 per group) were randomly assigned to receive daily oral doses of either vehicle (citric acid with sodium citrate monobasic) or the pan-RAS inhibitor ADT-1004 (40 mg/kg) until the conclusion of the experimental period. At the study’s end, mice were euthanized, and tumor images and weights were documented. The UAB’s IACUC approved the experimental protocol for these mouse studies.

#### Immunohistochemical Staining and Evaluation

Animal tumor tissues were fixed in 10% formalin for 24 h, dehydrated, paraffin embedded, and sectioned with 5µM thickness for immunohistochemistry. Antigen retrieval was performed from the sectioned slides by using 10 mM citrate buffer, pH 6.0 for 5 min (microwave) for all the antibodies used (phospho-ERK, α-SMA, and Ki67). The slides were then rinsed with PBS followed by blocking with 5% BSA for 30 min. The sections were then probed with the primary antibodies overnight at 4 °C as follows: αSMA (Abcam, #ab7817, 1:500), phospho-ERK (CST, #4370, 1:500), and Ki-67 (CST, #9449 1:500). Following the antibody incubation, the slides were washed and incubated for 30 min with biotinylated anti-mouse/rabbit IgG secondary antibody. Afterwards, the slides were washed and incubated with VectaStain Elite ABC reagent (Vector Laboratories, #PK-8200) for 30 min and the counterstaining was performed for 1 min with hematoxylin reagent (Sigma-Aldrich, #51275), followed by washing and mounting with mounting medium (epredia, #4112). Stained slides were imaged using the Echo Revolution automated microscope (ECHO, USA) at 20× magnification, and quantified using ImageJ with the same threshold for each stain and the results are shown as percent staining per visual field.

### Multiplex Immunofluorescence

#### Hyperplex staining and whole-slide imaging

FFPE (formalin fixed, paraffin embedded) slides containing vehicle and ADT-1004 treated TMAs (Tumor microarrays)were processed using Biogenex EZ Retriever system and EZ-AR2 Elegance (#HK546-XAK) at 107 °C for 15 min. Cyclical multiplex immunofluorescence was then performed using the Lunaphore COMET seqIF staining platform at the UAB Flow Cytometry and Single Cell core facility. In brief, all primary antibodies were diluted in multi-staining buffer (BU06, Lunaphore) **Table S1** and incubated for 4 min. Secondary antibodies diluted in antibody diluent (LI-COR # 927-65001) **Table S1** and DAPI counterstain were incubated for 2 min. Imaging was performed using an image buffer (BU09, Lunaphore) for the following exposure times: DAPI 80 ms, TRICTC 400 ms, and Cy5 200 ms. Elution after each cycle was performed for 2 min at 37 ^0^C in the elution buffer (BU07-L, Lunaphore) followed by a 30 sec quenching step with quenching buffer (BU08-L, Lunaphore). Upon completion of the experiment, an aligned, stitched, flat-field corrected 16-bit OME-TIFF image file was generated for each slide. Images were pre-processed to subtract autofluorescence from each channel using Lunaphore Horizon Viewer and exported for further analysis.

#### Image Segmentation and Quantification

Visiopharm Oncotopix Image Analysis suite was used to de-array the whole slide scans (allows the retention of each TMA core as an individual biological replicate), tissue regions hand selected to remove artifacts such as folds and debris, identify sub-tissue region of interests (epithelium vs other based on Keratin 17-19 stain thresholds), and cell segmentation (based on Nuclear Segmentation AI, Deep Learning app with Keratin 17-19, CD3, CD11b and PDGFRb stains for membrane boundaries). Cellular phenotyping was performed using the change by intensity post-processing tool output in an app for known marker populations in a stepwise fashion with the undefined cell population resulting in the following populations being also negative for the previous markers: Cancer –Epithelium (Ker17/19 high), Cancer dim (Ker 17/19 dim), B cells (Ker-17/19-CD19^+^), T cells (Ker-17/19-CD19-CD3^+^), Cytotoxic T cells (Ker-17/19-CD19-CD3^+^CD8^+^), T effector (Ker-17/19-CD19-CD3^+^ CD4^+^), T regulatory (Ker-17/19-CD19-CD3^+^CD4^+^Foxp3^+^ and Ker-17/19-CD19-CD3^+^Foxp3^+^ if CD4 membranous stain was outside of the sectioning plane), macrophages (Ker-17/19-CD19-CD3-CD11b^+^), neutrophils (Ker-17/19-CD19-CD3-CD11b-Ly6G^+^), myeloid derived suppressor cells (Ker-17/19-CD19-CD3-CD11b-CD11b^+^Ly6G^+^), endothelial (Ker-17/19-CD19-CD3-CD11b-CD11b-Ly6G-CD31^+^), and cancer associated fibroblasts (negative for all other markers and positive for any of the mesenchymal markers: αSMA, FAP, PDGFRβ, desmin and integrinβ3). Expression of functional markers whose expression was of interest regardless of cellular population (pAKT, pERK, CD11b, αSMA, FAP, PDGFRβ, Desmin and Integrinβ3) was exported using the phenoplex guided workflow. Cellular features such as phenotype, marker positivity, x, y coordinates, area, and shapes (form factor and ellipticalness) were exported for each cell. Count analyses and percentages were calculated in xcel using pivot tables and graphed in Prism.

#### Spatial Analysis

Cell-cell infiltration was calculated using the G-Function for each pairwise target and neighbor analysis every 10um radii for 10-200um for each TMA core. Vehicle and ADT-1004 treated tumor cores were averaged and significant differences at each radii point were calculated using a t-test.

Marker expression analysis of interesting co-localization phenotypes were calculated using a MATLAB script delineated as follows:

For cells type X, type Y and marker A

Under a given radius r, assume that we can find:

X1 cells for cells that are: X type, expressing A, and has at least one Y cell within r

X2 cells for cells that are: X type, not expressing A, and has at least one Y cell within r X3 cells for cells that are: X type, expressing A, and has no Y cells within r

X4 cells for cells that are: X type, not expressing A, and has no Y cells within r X5 cells that are: X type, has at least one A-positive Y cell within r

X6 cells that are: X type, has at least one A-negative Y cell within r X7 cells that are: X type, no A-positive Y cells within r

X8 cells that are: X type, no A-negative Y cells within r

X9 cells for cells that are: X type, has at least one Y cell within r (total number of X co-localization with Y)

Sum of X1∼X4 will be the total number of X cells. X1+X3 will be the total number of A-expressing X cells.

X1+X2 will be the total number of X cells with co-localization with r.

X5+X7 = X6+X8 = X = total number of X.

We then define:

M1 = X1/(X1+X2). M1 is the fraction of A-expression in X that co-localize with Y

M2 = X3/(X3+X4). M2 is the fraction of A-expression in X that do not colocalize with Y M3 = X1/(X1+X3).

M3 is the fraction of co-localization of X with Y out of all A-expressing X cells

M4 = X2/(X2+X4). M4 is the fraction of colocalized X cells out of all non-A-expressing X cells

M5 = X5/X. M5 is the fraction of X that co-localizes with A-positive Y M6 = X6/X.

M6 is the fraction of X that co-localizes with A-negative Y.

### Statistical Analyses

Statistical analyses and data visualization were done using GraphPad Prism (version 9.1.2). The data are represented as means accompanied by either standard deviation (SD) or standard error of the mean (SEM). A repeated measures analysis of variance (ANOVA) or ANOVA with Bonferroni correction and a two-tailed Student’s t-test were conducted to evaluate and apply multiple corrections for assessing statistical significance between groups. To determine IC_50_ values, concentrations were log-transformed, and the data were normalized to the control, followed by a log (inhibitor) vs. response (four-parameter nonlinear dose-response) analysis. A statistical significance threshold was set at p < 0.05.

## Results

### ADT-1004 inhibits tumor growth in murine models of PDAC

Initial experiments were conducted to confirm oral bioavailability and sufficient conversion of the prodrug, ADT-1004, to it active metaobolite, ADT-007 *in vivo.* Following administration of a single oral dose of 100 mg/kg ADT-1004 generated sustained plasma levels of ADT-007 for at least 8 h that were appreciably higher than growth IC_50_ values (**Figure 1B**). Next, the maximum tolerated dose (MTD) of ADT-1004 was established when administered by oral gavage BID, 5x/week for 3.5 weeks using tumor-bearing mice. The results revealed no significant differences in body weight between the vehicle and ADT-1004 treated groups at dosages up to 175 mg/kg BID (**Figure 1C**). Dosages above 175 mg/kg BID resulted in significant weight loss (data not shown). Blinded analysis of H&E stained tissues from mice treated with 175 mg/kg BID ADT-1004 confirmed no discernable histopathologic abnormalities in the heart, lungs, kidneys, liver, jejunum, pancreas, and colon (**Figure 1D**). In addition, analysis of clinical chemistries, complete blood counts, and bone marrow cytology revealed no significant differences in markers of liver and renal function, electrolytes, energy metabolism, and hematologic function between mice treated with vehicle or ADT-1004 at the 175 mg/kg BID dose (**Supplementary** Figure 1).

To determine antitumor activity of ADT-1004, luciferase-labeled KPC (KPC-Luc) cells with point mutations in KRAS^G12D^ and p53^R172H^ were surgically implanted into the pancreas of C57BL6/J mice and imaged one week after implantation (day 0) to randomize mice between the control (vehicle) and ADT-1004 treatment groups. Treatment was started on day 0 by once daily oral gavage with either vehicle or ADT-1004 at dosages ranging from 10 – 50 mg/kg (5x/week) (**Figure 1E**). Mice were euthanized on day 27 to determine treatment effects on tumor growth by measuring bioluminescence and tumor weight (**Figure 1F**). Most mice in the ADT-1004 treatment groups displayed reduced bioluminescence compared with mice treated with vehicle. Doses of ADT-1004 from 20-50 mg/kg demonstrated comparable statistically significant reductions in tumor weights compared to vehicle-treated mice (**Figure 1F**) with no significant differences in body weight between the treated and vehicle groups (**Figure 1G**). Measurement of ADT-1004 and ADT-007 levels in plasma by LC-MS/MS from ADT-1004 treated mice revealed that the 40 mg/kg dose generated comparable levels as a 50 mg/kg dosage (**Figure 1H** and **1I**). We therefore selected 40 mg/kg dose for further investigation as this dose appeared to be the lowest possible dose that could achieve maximum systemic levels of ADT-007.

### Tumor growth inhibition by ADT-1004 is associated with reduced activated RAS and pERK levels

The mechanism responsible for the anti-cancer activity of ADT-007 was previously reported to involve direct binding to RAS within the nucleotide-binding domain to block GTP activation of RAS and suppress effector interactions and downstream MAPK/AKT signaling.^15^ We therefore conducted a second experiment using the KPC orthotopic model (with KRAS^G12D^ mutation) described above to confirm that ADT-1004, as a prodrug of ADT-007, inhibits tumor growth by blocking RAS activation. Similar to results from the first experiment, oral administration of ADT-1004 at a dosage of 40 mg/kg demonstrated a significant reduction in tumor growth compared with the vehicle group, as evidenced by the bioluminescence imaging and tumor weight without differences in body weight (**Figure 2A-E**). The tumor tissues were analyzed for immune changes using multi-parameter flowcytometry to identify myeloid, T, NK, and tumor cells in the TME (**Supplemental Figure 8**). Consistent with *in vitro* studies of ADT-007,^15^ measurement of activated RAS levels by RAS-RBD pull-down assays showed that tumors from ADT-1004 treated mice had significantly reduced RAS-GTP levels compared with tumors from vehicle-treated mice (**Figure 2G**). Tumors from mice treated with ADT-1004 treatment group also showed reduced pERK levels compared with tumors from the vehicle group (**Figure 2H**).

**Figure 2:**
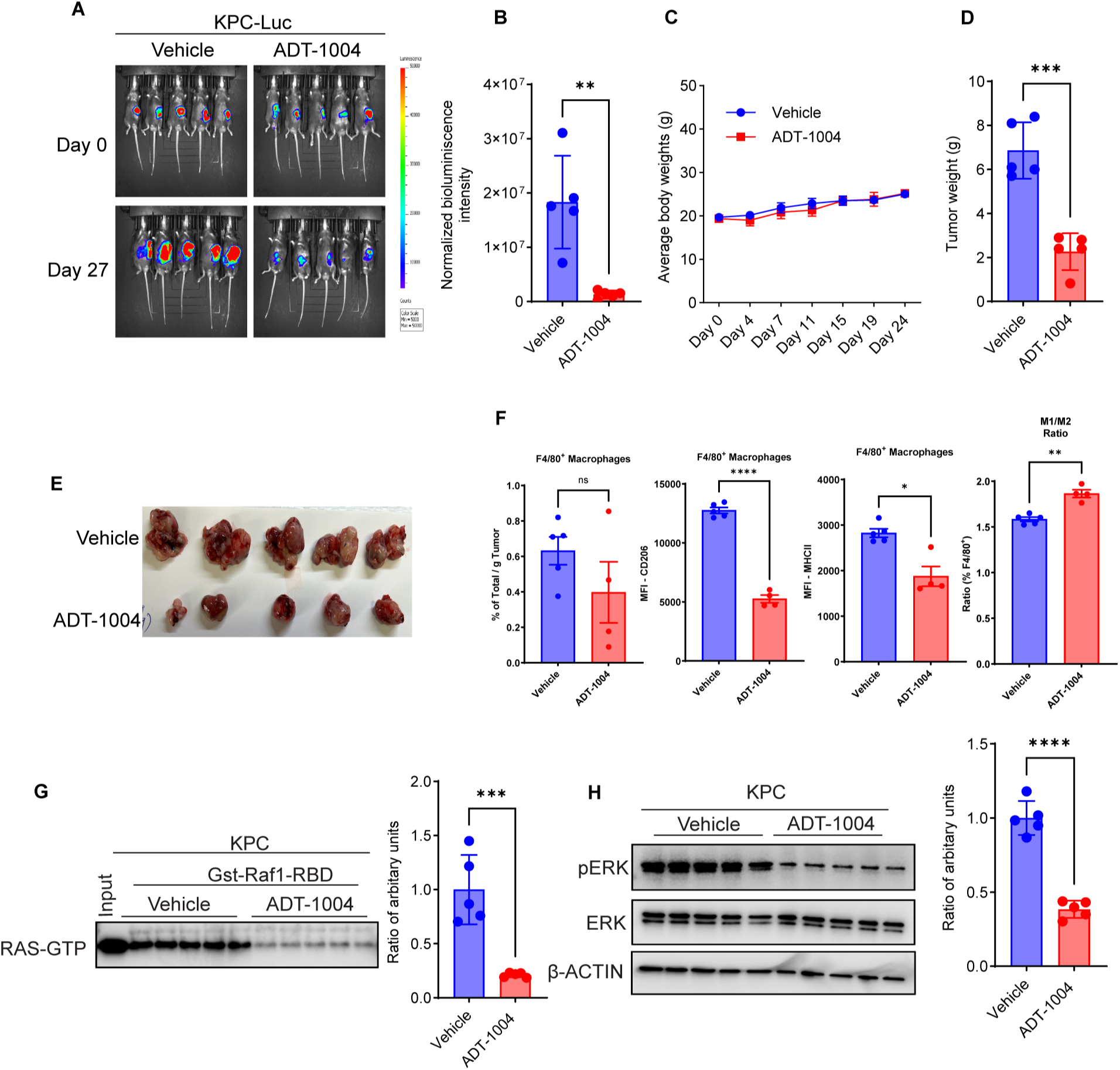
Effect of ADT-1004 treatment on tumor growth and molecular markers in KPC cell-implanted C57BL/6J mice. **(A).** KPC-Luc cells were injected orthotopically into the pancreas of C57BL/6J mice. Mice were treated by oral gavage 5days/week with vehicle or 40 mg/kg ADT-1004 and sacrificed after 27 days of treatment. Representative bioluminescence images at the indicated time points are shown. **(B).** Relative normalized whole-body bioluminescence intensities in mice under the indicated conditions (n=5). **(C).** Average body weights of mice treated with vehicle and ADT-1004 (40 mg/kg). **(D).** Tumor weights were measured from mice at the end of the experiment under the indicated conditions for experiment in (**A**). **(E).** Tumor images at the end of the experiment are shown. (**F**) The percentages of CD45^+^ CD11b^+^ F4/80, and CD206^+^ (macrophages), CD11b^+^ F4/80^+^ PD-L1^+^, and M1/M2 ratios increased in tumors from ADT-1004 treated (n=4) compared to vehicle (n=5) treated mice. Welch’s t-test determined p values. Error bars indicate SD. **(G).** Ithe tumors were probed for detecting activated RAS GTP levels by GST-Raf1-RBD pull-down assay (left) and graph depicting quantifyication of activated RAS GTP levels (right). **(H).** Indicated tumor tissues were analyzed for pERK and total ERK by immunoblotting. β-Actin was used as a loading control (left). Graph depicting quantifyication of pERK by western blot analysis, normalized using β-Actin (right). All quantitative data represent the mean ± SEM. ns, non-significant, ∗p < 0.05, ∗∗p < 0.01, ∗∗∗p < 0.001, and ∗∗∗∗p < 0.0001.

Next, we determined the antitumor activity of ADT-1004 in additional mouse PDAC models. First, we employed a different syngeneic, immunocompetent orthotopic mouse model in which 2838c3-luc cells were implanted into the pancreas of female C57BL6/J mice (**Figure 3A**). Mice were treated 5x/week with either vehicle or ADT-1004 at 20 mg/kg QD, 20 mg/kg BID, or 40 mg/kg QD. Measurement of bioluminescence and tumor weights showed that all three ADT-1004 dose groups displayed comparable significantly reduced tumor size compared with the vehicle group with no significant differences in body weight (**Figure 3A-E**).

**Figure 3:**
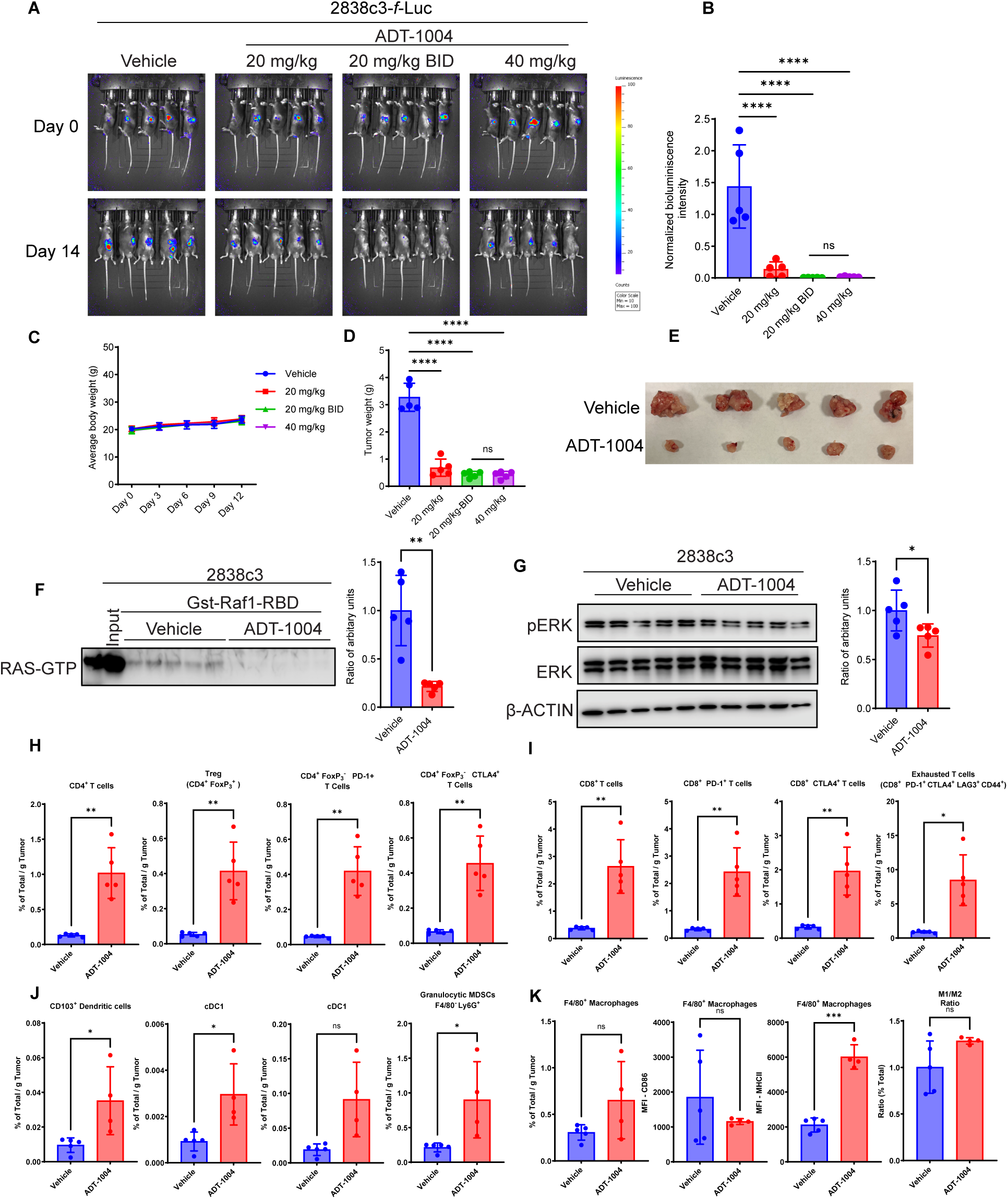
ADT-1004 treatment suppresses tumor growth and modulates tumor microenvironment in C57BL/6J mice with 2838c3 cell implants. **(A).** F-Luc-labeled 2838c3 cells were orthotopically implanted into C57BL6/J mice (n=20). Four groups of 5 mice were treated with vehicle or ADT-1004 by oral gavage (20 mg/kg, 20 mg/kg BID, and 40 mg/kg body weight) 5 days/week per week. **(B).** Normalized bioluminescence intensities of the tumors in the 4 groups of mice at day 14 are shown. **(C).** Average body weights of the animals during the course of the treatment shown in experiment (**A**). **(D)**. Tumor weights were measured from mice at the end of the experiment for indicated dosing schedules (**A**). (**E**). Tumor images at the end of the experiment are shown for vehicle and ADT-1004 (40 mg/kg). (**F**). Indicated tumor tissues were probed for detecting activated RAS GTP levels by GST-Raf1-RBD pull-down assay (left). Graph depicting the quantification of activated RAS GTP levels (right). (**G**). Indicated tumor tissues were analyzed for pERK and total ERK. β-Actin was used as a loading control (left). Graph depicting the quantification of pERK in western blot analysis, normalized using β-Actin (right). (**H**). Proportions of CD4^+^ T-cells, CD4^+^ Foxp3^+^, CD4^+^ PD-1^+^ T cells, and CD4^+^ CTLA-4^+^ T cells in 2838c3 TiME were determined by multi-parameter flow cytometry in ADT-1004 (n=5) and vehicle (n=5) treated mice. Welch’s unpaired t-test determined p values, ns indicates not significant. Error bars indicate SD. (**I**). Expression of CD8^+^ T-cells, CD8^+^ PD-1^+^ T cells, CD8^+^ PD-1^+^ LAG3^+^, and CD8^+^ PD-1^+^ CTLA-4^+^ subsets from 2838c3 TME. Welch’s t-test determined p values and error bars indicate SD. (**J**). The percentages of CD11b^+^ MHCII^hi^, XCR1^+^ cDC1, and CD11b^+^ CD11c^hi^ MHCII ^hi^ CD172α^+^ cDC2 subsets increased in ADT-1004 (n=5) compared to vehicle (n=5) treated mice. Welch’s t-test determined p values; ns indicates not significant. Error bars indicate SD. (**K**). The percentages of CD45^+^ CD11b^+^ F4/80, and CD206^+^ (macrophages), CD11b^+^ F4/80^+^ PD-L1^+^, M1/M2 ratio increased in ADT-1004 treated (n=5) compared to vehicle (n=5) treated mice. Welch’s t-test determined p values. Error bars indicate SD. ns, non-significant, ∗p < 0.05, ∗∗p < 0.01, ∗∗∗p < 0.001, and ∗∗∗∗p < 0.0001.

We also determined if the antitumor activity of ADT-1004 in the 2838c3 orthotopic PDAC model was associated with reduced levels of activated RAS. Similar to results in the KPC orthotopic model, RAS-RBD pull-down assays showed a significant reduction of RAS-GTP levels in tumors from ADT-1004-treated mice compared with tumors from vehicle-treated mice (**Figure 3F**). A notable decrease in the pERK levels was also observed in mice treated with ADT-1004 at 40 mg/kg, providing additional evidence that the antitumor activity of ADT-1004 is mediated by inhibition of activated RAS (**Figure 3G**) and consistent with previous *in vitro* studies of ADT-007.^15^

### ADT-1004 alters tumor immune microenvironment (TiME) by inhibiting KRAS^G12D^

Considering the growing literature reporting that mutation-specific KRAS inhibitors can enhance tumor-specific immunity,^18^ ^19^ ^20^ we investigated the activity of ADT-1004 in two PDAC-bearing, syngeneic, immunocompetent mouse models harboring the KRAS^G12D^ mutation. We selected the KRAS^G12D^ mutant luciferase expressing KPC and 2838c3 PDAC lines as both exhibit increased T cell infiltration and an effector phenotype.^15^ Moreover, the effector phenotype of CD8^+^ T cells as evidenced by high levels of PD-1 and CTLA4 expression was previously demonstrated in the 2838c3 model is responsive to combinatorial ICB (immune checkpoint blockade) utilizing antagonists to PD-1 and CTLA4.^21^ Given our previous findings involving direct injection of ADT-007 in subcutaneous syngeneic mouse PDAC models, including 2838c3,^15^ we assessed the numbers and phenotype of T, NK, and myeloid cell subsets by multiplarameter flow cytometry (**Supplementary** Figure 2) to determine whether ADT-1004, as a prodrug of ADT-007, had a similar impact on immune subset abundances and phenotypes in the pancreatic TME.

Following 28 days of treatment course (5days/week), mice bearing KPC or 2838c3 tumors were euthanized to evaluate changes in the immune cell population densities within the pancreatic TME. ADT-1004 treatment did not alter density (% Total/gm tumor) of innate-like (NKT and γδT) or NK cells in the KPC model (**Supplementary** Figure 2A), compared with the 2838c3 model where the densities of innate-like T and NK cell subsets were increased with treatment (**Supplementary** Figure 2B). Furthermore, ADT-1004 treatment increased the density of CD4^+^ and CD8^+^ T cells in the 2838c3 TME (**Figure 3H-I**) despite remaining unchanged in the KPC TME (**Supplementary** Figure 3A-B). Phenotypic analysis revealed that ADT-1004 treatment caused a marked increase in the density of CD4^+^ and CD8^+^ PD-1^+^ and CD4^+^ CTLA4^+^. (**Figure 3H-I**). Furthermore, the densities of CD4^+^ and CD8^+^ T cells co-expressing PD-1, CTLA4, LAG-3, and CD44 (**Figure 3I**) were increased in 2838c3 tumors after ADT-1004 treatment, indicating an emergence in exhaustion phenotype. Overall, these findings indicate an increase in the density of activated and potentially exhausted T cells after ADT-1004 treatment.

Analysis of myeloid cells after ADT-1004 treatment revealed an increased density (% total/mg tumor) of F4/80^+^ macrophages, XCR1^+^ cDC1, CD172α^+^ cDC2, and Ly6G^+^ granulocytes in the 2838c3, but not KPC TME (**Figure 3J+K and Supplementary** Figure 3C). In each model, F4/80^+^ macrophages exhibited an increase ratio of M1 like (MHCII^hi^) macrophages compared to M2 (CD206^+^) like macrophages (**Figure 2F and Figure 3K**). An increase in pro-inflammatory M1 macrophage and dendritic cell subsets is suggestive to a TME allowing for improved anti-tumor immune responses, and increased expression of immune checkpoint receptors PD-1, CTLA4, and LAG-3, indicating a potential benefit of combination with antagonists to these pathway with ADT-1004.

### Spatial phenotyping reveals enhancement of TME infiltration in ADT-1004 treated tumors

We next sought to investigate the effects of ADT-1004 on the cellular infiltration patterns o within the tumor core. Pancreatic tumors from the 2838c3-luc murine PDAC model were initially assessed for viable tumor cores by a licensed veterinary pathologist and 2 mm cores were obtained to generate a microarray of each tumor (n=4 for each treatment group). We performed multiplex immunofluorescence using the Lunaphore COMET system and a 15-plex antibody marker panel for general neoplastic, immune, and stromal cell profiling (**Figure 4A**). Cells were classified into 13 distinct phenotypes based on marker positivity: cancer cells CK^Bright^, cancer cells CK^Dim^, B-cells (CD19^+^), undefined T cells (Tcells (other): CD3^+^CD8^-^CD4^-^), cytotoxic T cells (CytoT: CD3^+^ CD8^+^), T effector cells (Teff: CD3^+^ CD4^+^), T regulatory cells (Treg: CD3^+^ CD4^+^ Foxp3^+^), macrophages (Mac: CD11b^+^ Ly6G^-^), Neutrophils (Neutro: CD11b^-^ Ly6G^+^), myeloid derived suppressor cells (MDSC: CD11b^+^ Ly6G^+^), cancer associated fibroblasts (CAF: any combination of FAP^+^, αSMA^+^, PDGFRβ^+^, Desmin^+^, or Integrinβ3^+^), and undefined cells (negative for all markers).

**Figure 4:**
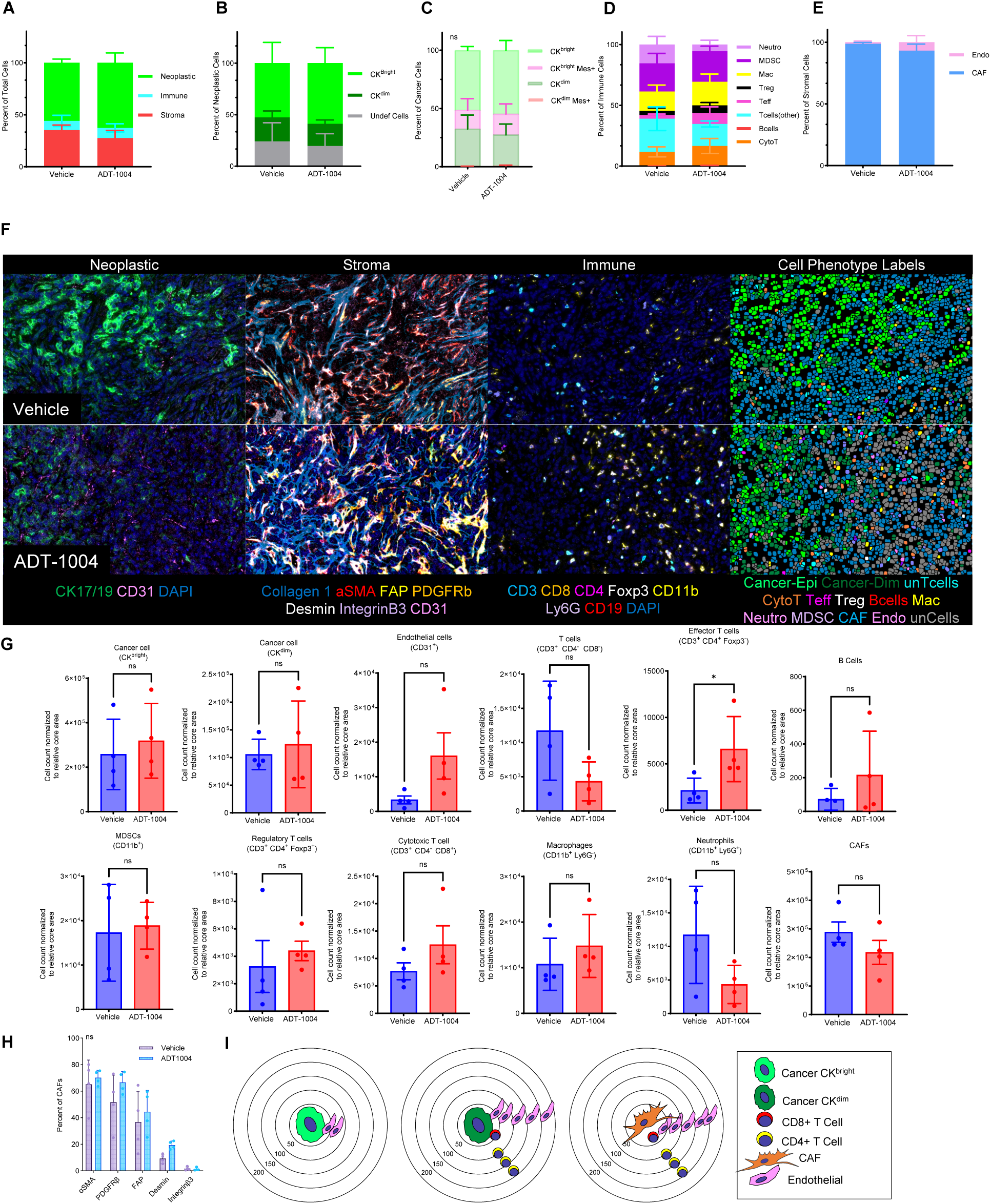
Spatial quantification reveals increased immune infiltration in ADT-1004 treated TME. **(A).** The expression matrix for each of the cell phenotypes based on Visiopharm results. **(B).** Percentage of neoplastic cells (y-axis) in each tumor (color legend) for each core of the TMA (x-axis). **(C).** Mesenchymal expressing CK^dim^ cells in ADT-1004 treated tumors did not co-localize with CD8+ or CD4+ T cells. **(D).** Percentage of immune cells (y-axis) in each tumor (color legend) for each core of the TMA (x-axis). **(E).** Annotations of classified neoplastic and immune cells (top) in the vehicle (left) and ADT-1004 treatments (right) and the corresponding antigens detected in the multi-color fluorescence images (bottom). **(F).** The t-distributed stochastic neighbor embedding (tSNE) plot showed 13 clusters (color legend) in a single core (left). Expression programs of different cell phenotypes (right) in the vehicle and ADT-1004 treated cores. tSNEs of single-nucleus profiles (dots) of neoplastic cells (cancer-epi, cancer-dim, and CAFs) from all tumors, and immune compartments (Macrophages, cyto T cells, T cells, Teff, and Tregs). **(G).** Unchanged CK^bright^ and CK^dim^ cells and increasing trend in endothelial cell infiltration in ADT-1004 treated TMAs compared to vehicle. Unchanged Tregs, significantly increased Teff, increased Teff/Treg ratio, and increased B cells in ADT-1004 treated TMAs compared to vehicle. Increased monocytic MDSCs (CD11b^+^ Ly6G^-^) and decreased granulocytic MDSCs (CD11b^+^ Ly6G^+^) in ADT-1004 treated TMAs compared to vehicle. Slightly reduced CAFs and increased endothelial cells in ADT-1004 treated TMAs compared to vehicle. **(H).** Unchanged percentage of CAFS in (α-SMA, PDGFRβ, FAP, Desmin, and Integrinβ3). **(I).** Reduced pericytic coverage in ADT-1004 treated tumors compared to vehicle in endothelial cells (left). Mesenchymal expressing CK^dim^ cells in ADT-1004 treated tumors did not co-localize with CD8^+^ or CD4^+^ T cells (middle). FAP^+^ and Integrinβ3^+^ CAFs exclude CD8^+^ T cells Integrinβ3^+^ CAFs exclude CD4^+^ T cells (right). ns: not significant and *p < 0.05.

Initially, count-based metrics showed no significant differences between the treatment groups in the proportion of neoplastic, stromal, or immune cell compartments (**Figure 4A**). Within the neoplastic cellular compartments, there were also no differences in the proportions of cancer cell subtypes, (**Figure 4B**) indicative of epithelial to mesenchymal transformation (EMT) status when assessing mesenchymal marker co-expression (**Figure 4C**). The immune cell counts showed a significant increase in T effector cells in the ADT-1004 treated tumors matching the observations obtained from the dissociative immune phenotyping (**Figure 3H-K**, **Figure 4D**). Within the stromal compartment there were also no significant differences in the percentages of CAFs both in aggregate and in the single marker expression patterns (**Figure 4E**). There was a trending increase in the number of endothelial and immune cell infiltrate in the ADT-1004 treated tumors (**Figure 4F**). These data show minimal differences in the cellular composition of the tumor core microenvironment upon ADT-1004 treatment.

To assess effects of ADT-1004 treatment on the cellular infiltration patterns of the core TME, we performed the G-function analysis which quantifies the infiltration of one cell-type (neighbor) within specified radii of another cell type (target). We performed this analysis for sets of 10 µm radii ranging from cell-cell contact distances 10-50 µm through non-cell contact distances (60-200 µm). Significance between vehicle and ADT-1004 were calculated for each pair-wise comparison at each radius. We observed both Cancer^CKbright^ and Cancer^CKdim^ cells had significantly more endothelial cell infiltration in the ADT-1004 treated tumors suggesting better vasculature coverage (**Figure 4G**). In addition to this, we have not identified any significant difference in the proportions of CAF percentage in ADT-1004 treated tumors compared too vehicle (**Figure 4I**). Both Cancer^CKdim^ and CAF cells had increased CytoT cell infiltration within cell-cell contact radii and Teff infiltration within near secretory distances of 100-150µm (**Figure 4I**, **Supplementary** Figure 4A-C), which correlate with known functions of cell-cell contact and cytokine secretion for these cell types.^22^

We next determined if specific mesenchymal markers (αSMA, FAP, Desmin, PDGFRβ, or Integrinβ3) in the target cells were associated with immune cell infiltration patterns. Cancer^CKdim^ cells that co-expressed any mesenchymal marker was not co-localized with the CytoT or Teff cells. This observation confirms previous spatial infiltration observations in mouse and human PDAC samples of reduced T cell infiltration around mesenchymal expressing cancer cells.^23^ Analysis of mesenchymal marker expression in the target CAF population showed FAP^+^ and Integrinβ3^+^ CAFs excluded CD8^+^ T cells in the ADT-1004 treated tumors and Integrinβ3^+^ CAFs excluded Teff cells; all other mesenchymal markers had no association with immune cell infiltration. Marker expression analysis of the neighbor cells showed that ADT-1004 treated tumors had a significant increase in infiltration of CD11b^-^ CytoT and Teff cells around Cancer^CKdim^ and CAF cells. These data demonstrate enhancement of immune cell infiltration with ADT-1004 treatment.

### ADT-1004 inhibits the growth of KRAS mutant PDX tumors

Given previous reports that ADT-007 inhibits the growth of KRAS mutantcancer cells regardless of the KRAS mutation, we determined the antitumor activity of ADT-1004 across a panel of three PDAC PDX models harboring different KRAS mutations (KRAS^G12D^, KRAS^G12C^, and KRAS^G13Q^) and one pancreaticobiliary PDX model (KRAS^G12V^). Tumor fragments from each of the PDX models were subcutaneously implanted into the right flank of NSG mice. Body weight and tumor volumes were measured while the mice were treated orally with ADT-1004 once daily at a dose of 40 mg/kg (5days/week). Tumor weights were determined at the end of the experiment. ADT-1004 treatment significantly inhibited the growth of PDAC tumors with KRAS^G12D^ (**Figure 5A and 5C, Supplemental Figure 5A**), KRAS^G12C^ (**Figure 5E and 5G, Supplemental Figure 5B**), KRAS^G12V^ (**Figure 5I and 5K, Supplemental Figure 5C**) and KRAS^G13Q^ (**Figure 5M and 5O, Supplemental Figure 5D**) with no change in body weight over the course of treatment, further demonstrating tolerability at dosages that inhibit tumor growth (**Figure 5B, 5F, 5J**, and **5N**, respectively) and therefore the potential for clinical development of ADT-1004 for PDAC.

**Figure 5:**
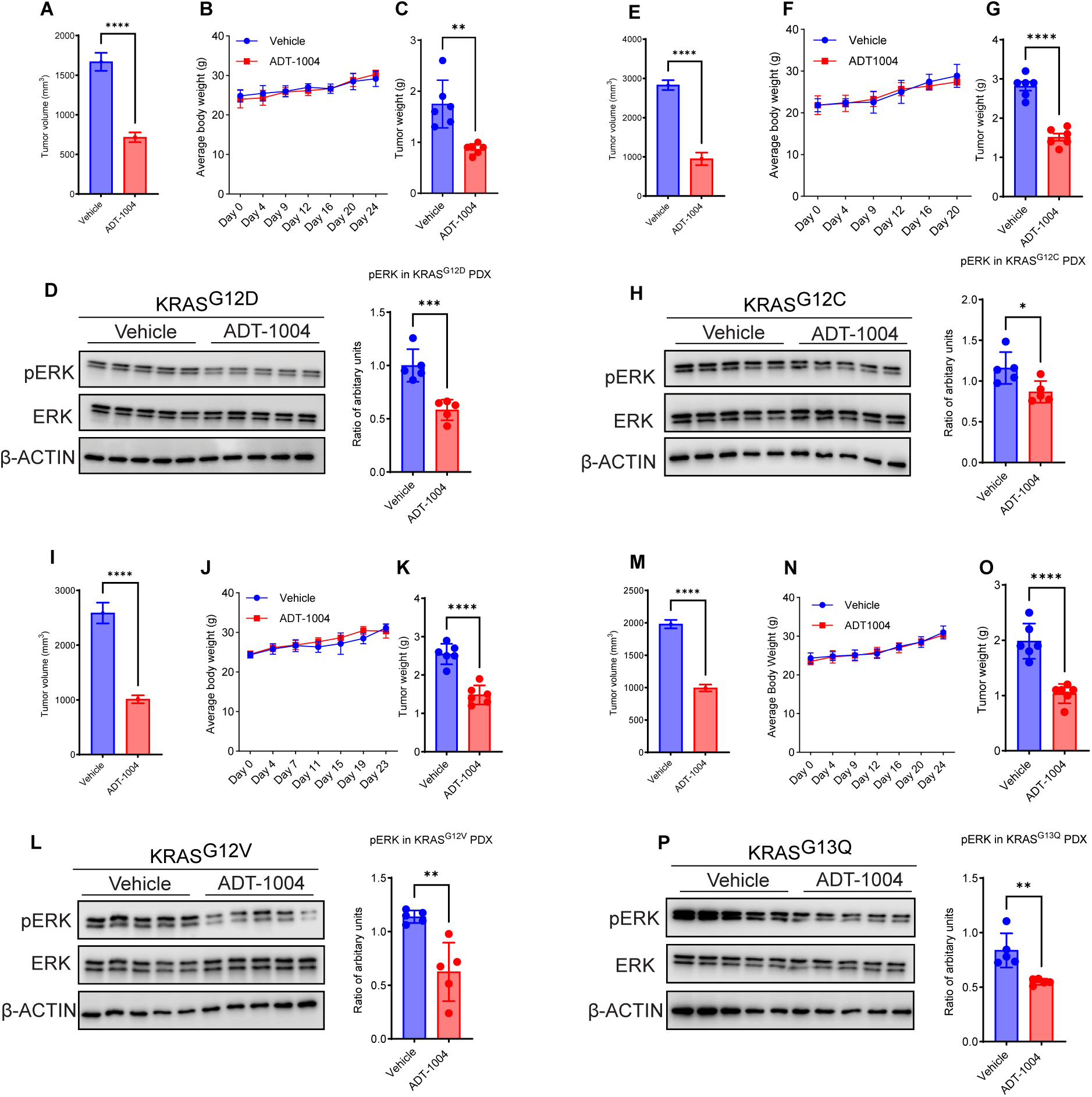
ADT-1004 treatment suppresses tumor growth and downregulates pERK levels in KRAS-mutant PDX models. (**A-D**). KRAS^G12D^ PDX. (**A**) Tumor growth curves of KRAS^G12D^ PDX implanted subcutaneously into NSG mice and treated with vehicle or ADT-1004 (n=6). (**B**). Average body weights of mice treated with vehicle and ADT-1004 (40 mg/kg). (**C**). Tumor weights were measured from mice at the end of the experiment under the indicated conditions for experiment in (**A**). (**D**). The indicated tumor tissues were probed for measuring pERK and total ERK. β-Actin was used as a loading control. Graph depicting the quantification of pERK by western blot analysis, normalized using β-Actin (left). (**E-H**) KRAS^G12C^ PDX. (**E**) Tumor growth curves of KRAS^G12C^ PDX implanted subcutaneously into NSG mice (n=6). (**F**). Body weights of mice treated with vehicle and ADT-1004 (40 mg/kg). (**G**). Tumor weights were measured from mice at the end of the experiment under the indicated conditions for experiment in (**E**). (**H**). The indicated tumor tissues were probed for measuring pERK and total ERK. β-Actin was used as loading control (right). Graph depicting the quantification of pERK by western blot analysis, normalized using β-Actin (left). (**I-L**). KRAS^G12C^ PDX. (**I**). Tumor growth curves of KRAS^G12V^ PDX implanted subcutaneously into NSG mice (n=6). (**J**). Average body weights of the animals during the course of the treatment shown in the experiment. (**K**). Tumor weights were measured from mice at the end of the experiment under the indicated conditions for experiment in (**I**). (**L**). The indicated tumor tissues were probed for measuring pERK and total ERK. β-Actin was used as loading control (right). Graph depicting the quantification of pERK by western blot analysis, normalized using β-Actin (left). (**M-P**) KRAS^G13Q^ PDX. (**M**)Tumor growth curves of KRAS^G13Q^ PDX implanted subcutaneously into NSG mice (n=6). (**N**). The average body weights of the animals during the treatment are shown in the experiment. (**O**) Tumor weights were measured from mice at the end of the experiment under the indicated conditions for experiment in (**M**). (**P**) The indicated tumor tissues were probed for measuring pERK and total ERK. β-Actin was used as loading control (right). Graph depicting the quantification of pERK by western blot analysis, normalized using β-Actin (left). Data represent the mean ± SEM. *p < 0.05, **p < 0.01, ***p < 0.001, and ****p < 0.0001.

Immunohistochemical (IHC) analysis of four KRAS mutant PDX tumors revealed the multiple effects of ADT-1004 treatment. We have identified a significant reduction of pERK levels in tumors from mice treated with ADT-1004 compared to mice treated with vehicle. In addition, αSMA immunostaining resulted in decreased expression within the ADT-1004 treatment group with distinct changes indicating structural reorganization, tissue remodeling, or fibrotic processes. Finally, analysis of Ki-67 levels showed a marked decrease in proliferative activity in all four treatment groups relative to their respective vehicle group, G12D (**Figure 6A**), G12C (**Figure 6B**), G12V (**Figure 6C**) and G13Q (**Figure 6D**) mutant expressing tumors.

**Figure 6:**
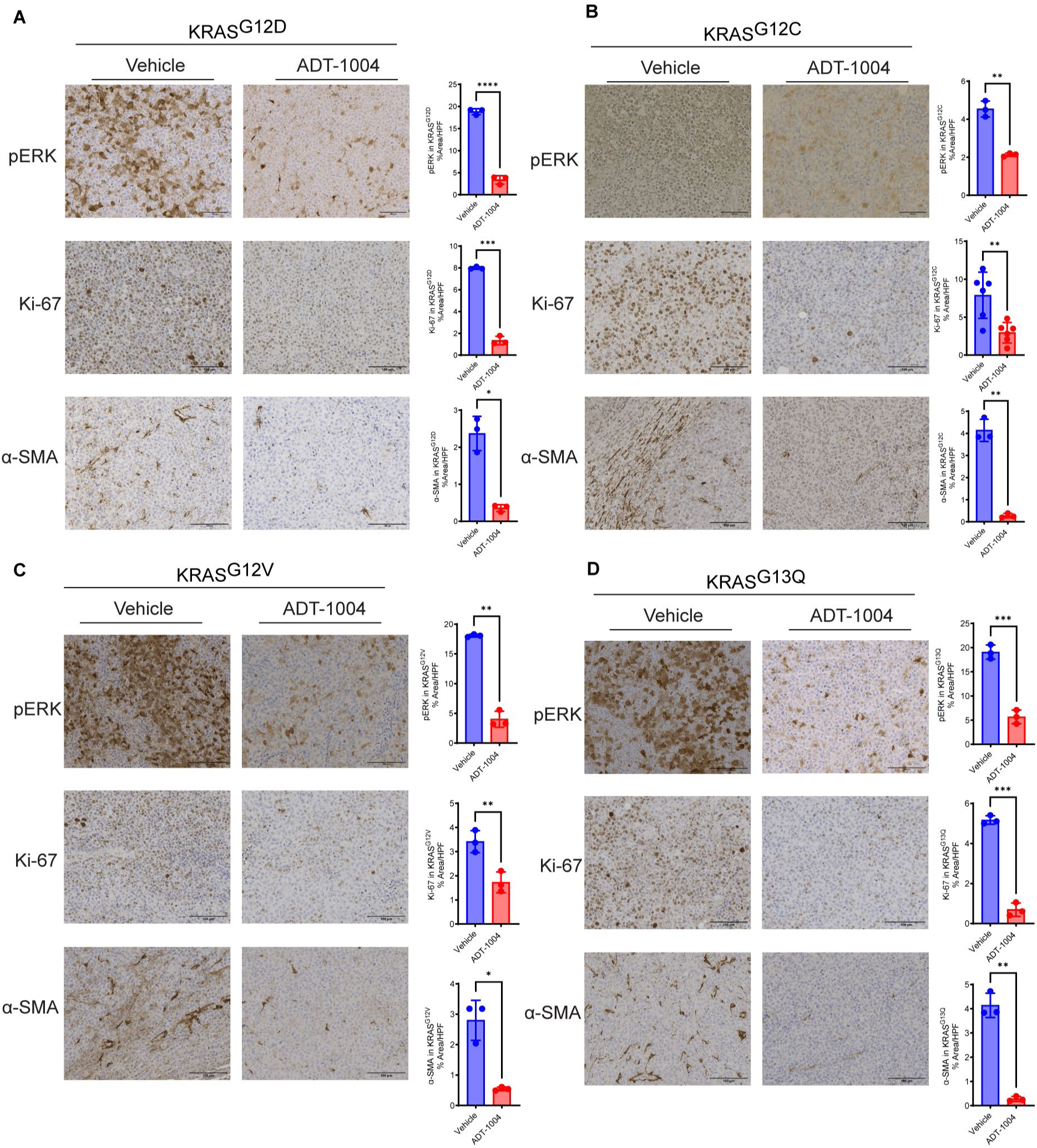
ADT-1004 treatment downregulates molecular markers in KRAS-mutant PDX models. (**A**) Representative IHC images of pERK, αSMA, and Ki-67 (left) and quantifications of IHC results (right) in KRAS^G12D^ PDX model. (**B**). Representative IHC images of pERK, αSMA, and Ki-67 (left) and quantifications of IHC results (right) in KRAS^G12C^ PDX model. (**C**). Representative IHC images of pERK, αSMA, and Ki-67 (left) and quantifications of IHC results (right) in KRAS^G12V^ PDX model. (**D**). Representative IHC images of pERK, αSMA, and Ki-67 (left) and quantifications of IHC results (right) in KRAS^G13Q^ PDX model. Results reveal that these proteins are highly expressed in KRAS mutations (G12D, G12V, G12C, and G13Q) and reduced expression in ADT-1004 treated tumors. Data represent the mean ± SEM. *p < 0.05, **p < 0.01, ***p < 0.001, and ****p < 0.0001.

### ADT-1004 does not inhibit tumor growth of RAS^WT^ PDAC cells

Additional experiments were performed to confirm the RAS selectivity of ADT-1004, given previous studies reporting that RAS^WT^ cancer cells with downstream BRAF mutations were einsensitive to ADT-007.^15^ Male NSG mice were implanted subcutaneously with BxPC-3 (RAS^WT^ BRAF^V600E^) cells and treated orally with 40 mg/kg ADT-1004 for 31 days. Body weight and tumor volume were monitored twice weekly throughout the study. ADT-1004 treatment did not affect BxPC-3 tumor growth (**Supplementary** Figure 6A) or impacted body weight (**Supplementary** Figure 6B), with no statistically significant difference observed in tumor weights compared to vehicle-treated mice (**Supplementary** Figure 6C-6D). These results lead us to conclude that ADT-1004 inhibits tumor growth as a highly selective pan-RAS inhibitor.

### ADT-1004 inhibits tumor growth of PDAC cells resistant to KRAS^G12C^ inhibitors

Human MIA PaCa-2 KRAS^G12C^ cells resistant to KRAS^G12C^ inhibitors (MIA-AMG-Res) were developed as previously described.^16^ Resistance to sotorasib and adagrasib was intitally confirmed by *in vitro* experiments to determine growth IC_50_ values against the parental MIA PaCa-2 cells. MIA PaCa-2 cells showed remarkable sensitivity to ADT-007 with an IC_50_ value of 3 nM, while sotorasib and adagrasib were less potent with IC_50_ values of 28 and 21 nM, respectively, and caused less overall growth inhibition (**Figure 7A**). MIA-AMG-Res cells were only slightly less sensitive to ADT-007 with an IC_50_ value of 16 nM, but completely lost sensitivity to sotorasib and adagrasib (**Figure 7B**). In addition, ADT-007, sotorasib, and adagrasib inhibited colony formation in parental MIA PaCa2 cells. However, the differences in potency and extent of killing caused by ADT-007 compared with sotorasib and adagrasib were more pronounced, which may reflect the ability of ADT-007 to induce mitotic arrest and apoptosis as previously reported.^15^ Similar to growth assays, ADT-007 retained activity to inhibit colony formation in MIA-AMG-Res cells, while sensitivity to sotorasib and adagrasib was markedly diminished (**Figure 7C** and **7D**).

**Figure 7:**
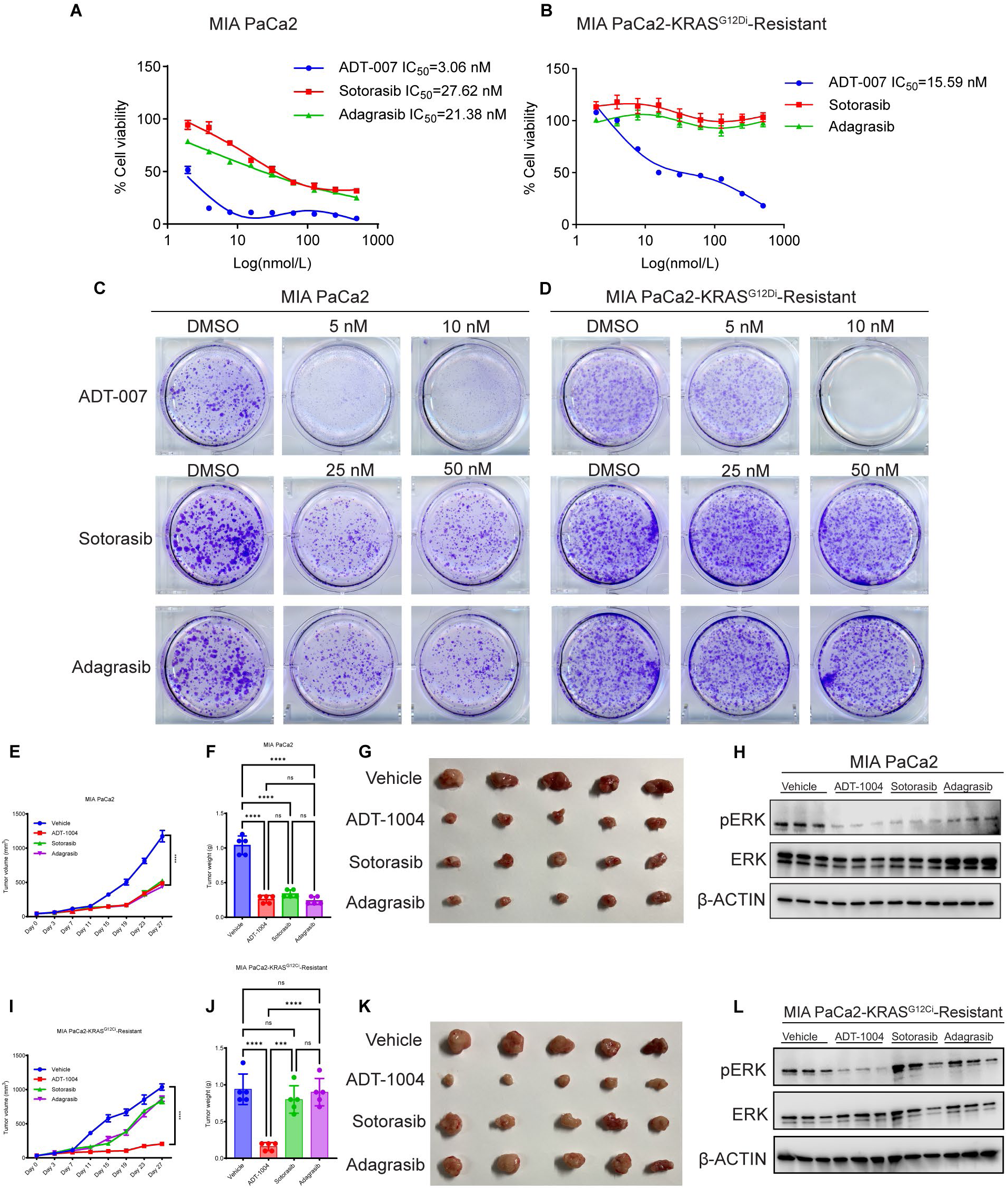
ADT-1004 demonstrates superior efficacy in suppressing the growth of KRAS^G12Ci-^resistant cells compared to AMG-510 and MRTX849. (**A-B**) PDAC cell lines were treated with different concentrations of ADT-007, sotorasib, and adagrasib for 3 days and analyzed for survival using the MTT assay. (**C-D**) Indicated PDAC cell lines were treated with the different concentrations of ADT-007, Sotorasib or Adagrasib for 2 weeks. Cell survival was then measured using clonogenic assays. Representative images are shown. (**E** and **I**) MIA PaCa-2 or MIA-AMG-Res cells were injected subcutaneously into the flanks of NSG mice (n=5). The mice were treated with either vehicle or ADT-1004, Sotorasib or Adagrasib (40 mg/Kg body weight) orally, 5 days/week for 4 weeks. The average tumor volumes over the course of the experiments are plotted. (**F** and **J**) Tumor weights were measured from mice in the vehicle ADT-1004, sotorasib, and adagrasib (40 mg/kg) groups at the end of the experiment under the indicated conditions for experiments in (**E** and **I**). (**G** and **K**) Representative tumor images of NSG mice in the vehicle, ADT-1004, Sotorasib, and Adagrasib (40 mg/kg) groups at the end of the experiment. (**H** and **L**) The indicated tumor tissues were probed for pERK and total ERK expression by immunoblotting. β-Actin was used as a loading control. Data represent the mean ± SEM. ns: not significant, *p < 0.05, **p < 0.01, ***p < 0.001, and ****p < 0.0001.

Next, experiments utilizing NSG mice xenografted subcutaneously with parental MIA PaCa-2 and MIA-AMG-Res cells were conducted. Parental MIA PaCa-2 cells showed comparable decreased tumor volume (**Figure 7E**) and final tumor weights (**Figure 7F** and **7G**), following treatment with ADT-1004, sotorasib or adagrasib in which each were tested at a once-daily dose of 40 mg/kg (5days/week). No significant differences in body weight were measured across treatment and vehicle groups (**Supplementary** Figure 7A). In addition, we found a significant reduction in pERK levels in tumor tissues in all the treatment groups compared to vehicle-treated mice tumor tissues (**Figure 7H** and **Supplementary** Figure 7C). In a side-by-side comparision, oral administration of ADT-1004 led to a dramatic decrease in tumor volume (**Figure 7I**) and weight (**Figure 7J** and **7K**) compared with vehicle against tumors derived from MIA-AMG-Res cells. By contrast, neither sotorasib nor adagrasib were effective. Notably, ADT-1004 demonstrated antitumor activity without any visible change in the body weights of treated mice (**Supplementary** Figure 7B). Furthermore, western blotting for the detection of pERK showed a significant reduction in the ADT-1004 treated tumors compared to all other treatment or vehicle groups, while meither sotorasib or adagrasib impacted pERK levels (**Figure 7L** and **Supplementary** Figure 7D). These results show that ADT-1004 is highly effective against KRAS^G12C^ tumors resistant to sotorasib or adagrasib, suggesting the potential to treat patiesnts who fail therapy with KRAS^G12C^ inhibitors.

## Discussion

PDAC poses a formidable global health challenge as the most lethal cancer with a notable absence of effective treatments.^24^ Point mutations in the RAS oncogene family, encompassing KRAS, NRAS, and HRAS are widespread across multiple cancers, but especially mutations in KRAS as a driver of PDAC. Despite the high prevalence of KRAS mutations in the most fatal human cancers, direct targeting of KRAS by reversible inhibitors has been challenging primarily because of the lack of binding surfaces on the protein and high affinity to bind GTP to inhibit GTP activation of effector interactions. The recent FDA approval of covalent KRAS^G12C^ inhibitors for the treatment of non-small cell lung cancer has renewed optimism to study RAS as a druggable target, although currently available KRAS^G12C^ inhibitors have limited use for PDAC given that KRAS^G12C^ mutations are rare in patients diagnosed with PDAC. Recognizing the role of other KRAS mutations in PDAC and other cancers, namely KRAS^G12D^, intensive efforts are ongoing to develop KRAS^G12D^ inhibitors.^25^ However, given that adaptive resistance is a crucial limitation of KRAS^G12C^ inhibitors that is attributed to various compensatory mechanisms, for example, from new KRAS mutations or activation of RAS^WT^ isoforms, likely other allele-specific inhibitors will have similar limitations.^14^

This present study builds upon our discovery of ADT-007, a reversible, highly potent, and selective pan-RAS inhibitor, with broad growth inhibitory activity and potential to avoid resistance mechanisms limiting the efficacy of mutant or isoform specific RAS inhibitors.^15^ As ADT-007 has low water solubility and is rapidly metabolized by glucuronidation, which limits systemic exposure by systemic IV or oral administration, we developed ADT-1004 as an orally bioavailable prodrug of ADT-007. Oral administration of ADT-1004 in mice generated sustained plasma levels of ADT-007 that was amenable to once or twice-daily dosing schedules, which produced appreciably higher ADT-007 concentrations than required to inhibit PDAC cell growth *in vitro*. The difference between the maximum tolerated dose of 175 mg/kg ADT-1004 twice-daily and the minimally effective dose of 40 mg/kg once-daily is relatively large, suggesting that ADT-1004 will have a broad therapeutic window in the clinic.

Here we showed antitumor activity of orally administered ADT-1004 against multiple mouse models of PDAC, including orthotopically implanted mouse cell lines and subcutaneously implanted PDX with different KRAS mutations. In these models, we consistently observed antitumor activity of ADT-1004 at a once-daily dose of 40mg/kg. Although higher dosages or a more frequent dosing schedule may result in greater antitumor activity, we selected 40 mg/kg as a minimal dosage that appeared to generate maximum plasma levels of ADT-007. In addition, this dosage demonstrated strong inhibitory effects on RAS activation and downstream signaling by measurement of RAS-GTP and pERK levels, respectively. These findings are also consistent with the published activity of other mutation specific and pan-Ras inhibitors in PDAC. ^26, 27, 28, 29^, although further studies are needed to determine which mechanism of inhibiting RAS will result in greater efficacy, tolerability, and the greatest potential to avert resistance.

A major challenge in using immune therapy for the treatment of PDAC has been the level of dysregulation of the immune system at multiple levels, including low T cells, exhausted T cells, absent antigen presenting cells and high inhibitory tumor-associated macrophages. Oncogenic KRAS is a significant enabler of the immune suppressive TME in PDAC such that blockage of RAS signaling promotes cytotoxic and M1 pro-inflammatory anti-tumor immune responses.^30^ Preclinical studies of mutant specific KRAS inhibitors, demonstrates a critical role of oncogenic KRAS signaling in immune suppression whereby targeting RAS vastly improves tumor immune responses.^31^ Our study demonstrates similar immune modulatory potential of ADT-1004 in comparison with other mutation-specific and pan-RAS inhibitors.^18, 20, 29, 32^ Specifically, treatment with ADT-1004 increases immune infiltration into the TME with increases in T cells infiltrations. Furthermore, there is an increase in T cells expressing immune suppressive receptors PD-1, CTLA4, and LAG-3, which are critical for immunosuppression of anti-tumor specific T cell responses.^33, 34^ Others have demonstrated that targeted inhibition of oncogenic KRAS^G12D^ signaling with MRTX1133 in pre-clinical models of PDAC resulted in increases in PD-1, such that blockade of this signaling pathway enhanced antitumor efficacy.^18,32^ Therefore, it is likely that ADT-1004 could enhance efficacy in comibination with agonists to PD-1 and CTLA4, although further studies are needed to assess this possibility along with tolerability. The observed increase in macrophage populations indicate a role for ADT-1004 in alteration of the macrophage phenotype from an immunosuppressive anti-inflammatory (M2 like) to a pro-inflammatory (M1 like) phenotype, which is again consistent with findings using targeted inhibitors. Additionally, we observed increases in the abundance of dendritic cell subsets, which are critical for priming tumor-antigen specific T cells.^35^ Interestingly CD11b^+^ Ly6G^+^ granulocytes in the 2838c3 TME were increased with ADT-1004 treatement. These cells expressed a significant, but modest increase in MHCII and downregulation of PD-L1, suggesting that these cells may exhibit antigen-presentation potential, which has been associated with increased survival in patients with a range of cancers and highlight another novel property of ADT-1004 to impact.^36^ Overall, this broad activation of macrophages (and granulocytes) with enhanced T cell priming capabilities suggests the potential of ADT-1004 to enhance antigen presentation and pro-inflammatory responses in the pancreatic TME.

Our spatial analysis of tumor core regions shows cell localization patterns were enhanced by ADT-1004 treatment. Of interest, is the enhancement of spatial patterns known in cancer biology as the paucity of infiltration of immune cells around mesenchymal-transitioned cancer cells and FAP^+^ and Integrinβ3^+^ CAFs. Combined with the observations of cancer cell line specific differences in immunological phenotypes, these observations suggest that tumor growth suppression by ADT-1004 treatment impacts host response mechanisms that are cancer specific. This activity also opens the possibilities for future combination therapies with TME targeting approaches.

Another desirable feature of ADT-1004 is its cytotoxic ability towards PDAC cells resistant to FDA-approved mutant-specific KRAS inhibitors. In this study, we show that human MIA PaCa-2 PDAC cells harboring KRAS^G12C^ developed to be resistant to sotorasib and adagrasib retained sensitivity to ADT-007 *in vitro* and to ADT-1004 *in vivo*. By contrast, both sotorasib and adagrasib essentially lost activity *in vitro* as determined by growth and colony formation assays and *in vivo* using a subcutaneous mouse tumor model. Previous clinical studies with KRAS^G12C^ inhibitors have identified multiple mechanisms of intrinsic or acquired resistance. These include new mutations in KRAS or activation of the wild-type KRAS allele or compensation from RAS WT isozymes (e.g., NRAS or HRAS) that are co-expressed with mutant KRAS and activated by growth factors (e.g., EGF) enriched in the TME.^14^ These resistance mechanisms can be bypassed by a pan-RAS inhibitor such as ADT-1004 and provide a key advantage over existing mutant-specific KRAS inhibitors to support clinical development. The ability of ADT-007 to induce mitotic arrest and apoptosis may provide additional advantages to avert resistance as previously reported.^15^ For example, ADT-007 completely inhibited colony formation of KRAS^G12C^ PDAC cells, while KRAS^G12C^ inhibitors (sotorasib and adagrasib), KRAS-specific inhibitors (BI-2865) and another pan-RAS inhibitor in development (RMC-6236) were appreciably less effective.

This comprehensive study establishes ADT-1004 as a 1^st^-in-class orally bioavailable pan-RAS inhibitor with promising antitumor efficacy against KRAS-mutant PDAC, including resistant phenotypes, which could be considered as a monotherapy. The changes in the TME observed with ADT-1004 also suggest that a combination with immune checkpoint inhibitors is a rational approach to evaluate in preclinical models.

