## Supplementary Figures for "ADT-1004: A First-in-Class, Orally Bioavailable Selective pan-RAS Inhibitor for Pancreatic Ductal Adenocarcinoma"

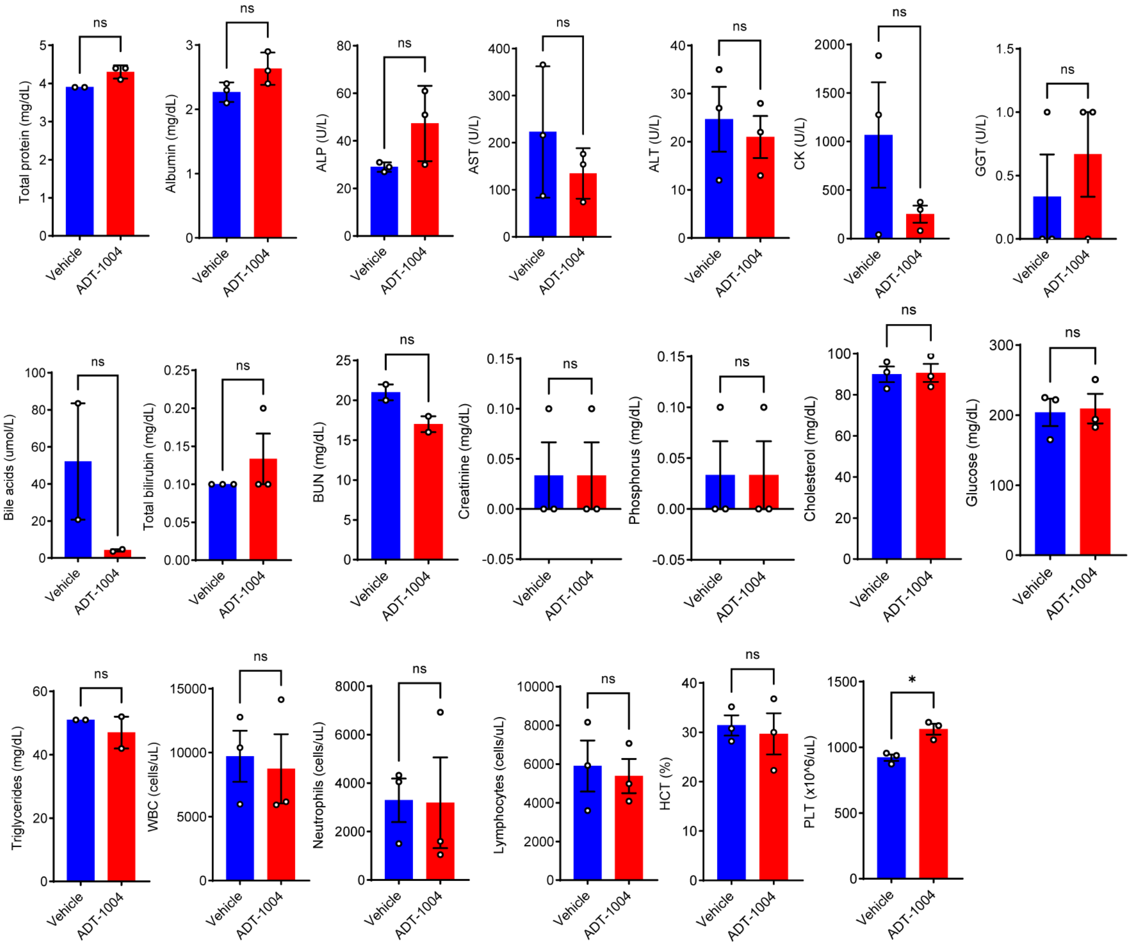


**Supplementary Figure 1: Serum biochemical analysis of mice treated with ADT-1004 (175 mg/kg BID).** Female C57BL/6 mice were implanted subcutaneously with 2838c3 PDAC cells and treated with ADT-1004 (175 mg/kg) orally, BID 5 days/week for 3.5 weeks. Serum was collected at the end of the treatment (n=3). Biochemical analysis indicated unchanged total protein, albumin, ALP, AST, ALT, CK, GGT, bile acids, total bilirubin, BUN, creatinine, phosphorus, cholesterol, glucose, triglycerides, WBC, neutrophils, lymphocytes, HCT and increased PLT as a result of ADT-1004 treatment compared to vehicle treatment (n=2-3/ test). ns: not significant, *: $p < 0.05$.

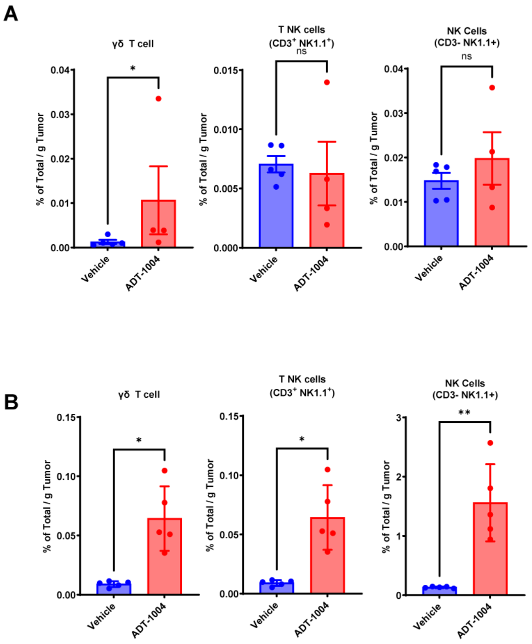


**Supplementary Figure 2. ADT-1004 treatment increases innate-like T and NK cells densities in the 2838c3 TiME. (A)** The density (% total/g tumor) of γδ T ($CD3^+$ $NK1.1^-$ TCR $\gamma\delta^+$), NKT ($CD3^+$ $NK1.1^+$TCR $\gamma\delta^-$), and NK ($CD3^-$ $NK1.1^+$ TCR $\gamma\delta^-$)cells were unchanged in the KPC TiME. **(B)** In comparison the densities (% total/g tumor) of these subsets were all significantly increased in the 2838c3 TME post ADT-1004 treatment. Data are representative of a single experiment of 5 mice/group. Cell densities were calculated by dividing the % total/tumor weight (g). Statistical significance was determined using Welch's t test. ns, non-significant, $*p < 0.05$, and $**p < 0.01$.

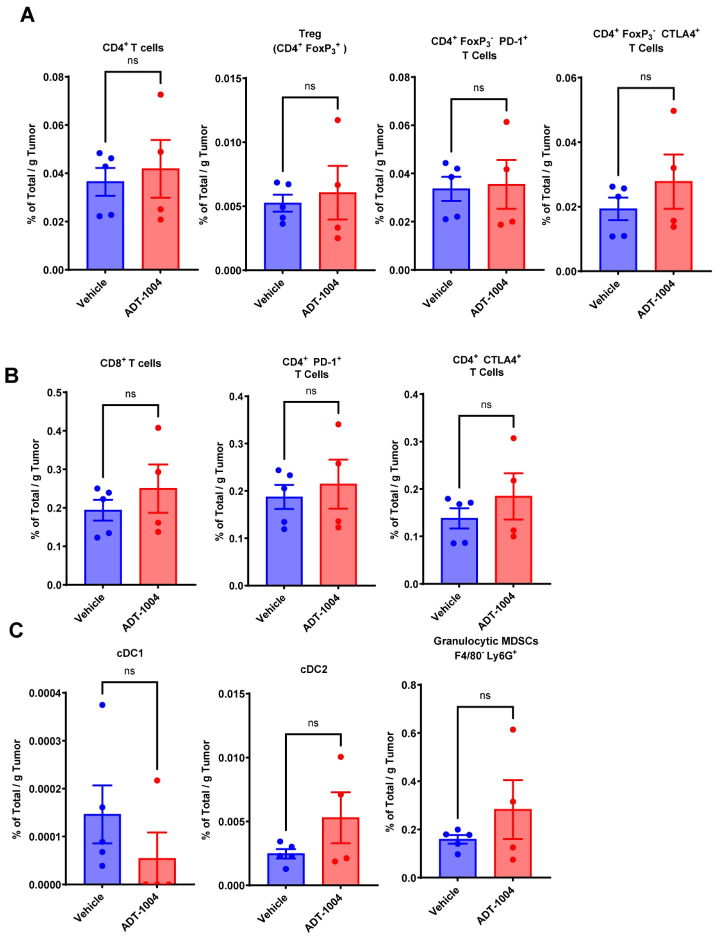


**Supplementary Figure 3. ADT-1004 treatment did not significantly alter density or phenotype of T cells in the KPC TME. (A)** Proportions of CD4$^+$ T-cells, CD4$^+$ Foxp3$^+$, CD4$^+$ PD-1$^+$ T cells, and CD4$^+$ CTLA-4$^+$ T cells in 2838c3 TiME were determined by multi-parameter flow cytometry in ADT-1004 (n=5) and vehicle (n=5) treated mice. Welch's unpaired t-test determined p values, ns indicates not significant. Error bars indicate SD. (**B**). Expression of CD8$^+$ T-cells, CD8$^+$ PD-1$^+$ T cells, CD8$^+$ PD-1$^+$ LAG3$^+$, and CD8$^+$ PD-1$^+$ CTLA-4$^+$ subsets from 2838c3 TME was graphed. Welch's t-test determined p values. Error bars indicate SD. (**C**). The percentages of CD11b$^+$ MHCII$^{hi}$, XCR1$^+$ cDC1, and CD11b$^+$ CD11c$^{hi}$ MHCII $^{hi}$ CD172α$^+$ cDC2 subsets increased in ADT-1004 (n=5) compared to vehicle (n=5) treated mice. Welch's t-test determined p values ns indicates not significant. ns, non-significant, $*p < 0.05$, $**p < 0.01$, $***p < 0.001$, and $****p < 0.0001$.

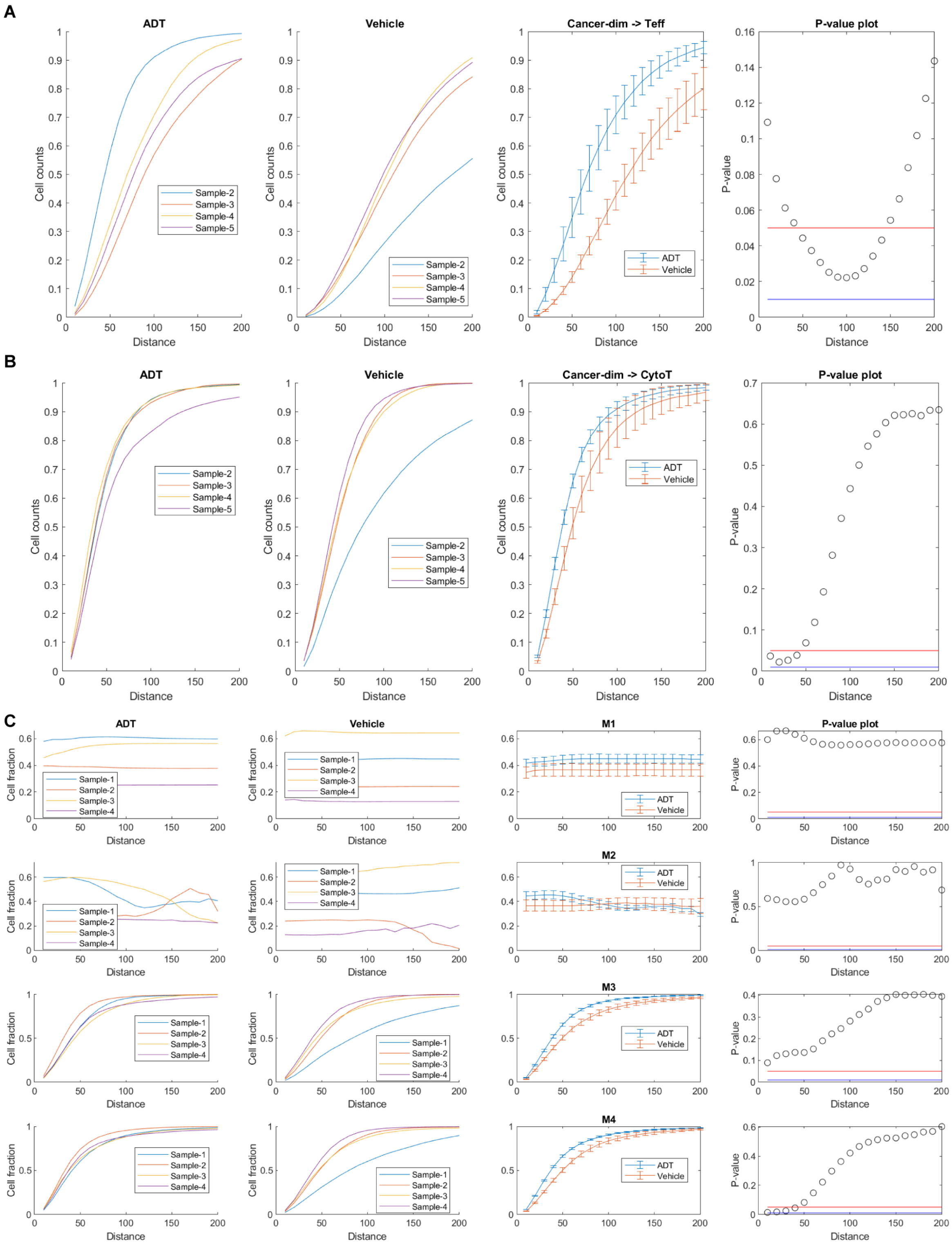


**Supplementary Figure 4: (A-B).** Spatial analysis within cell-cell contact radii showing increased CytoT cell infiltration near secretory distances of 100-150μm in both Cancer CK$^{dim}$ and CAFs. **(C).** Co-expression in immune cell infiltration patterns in CAF-FAP-CytoT population.

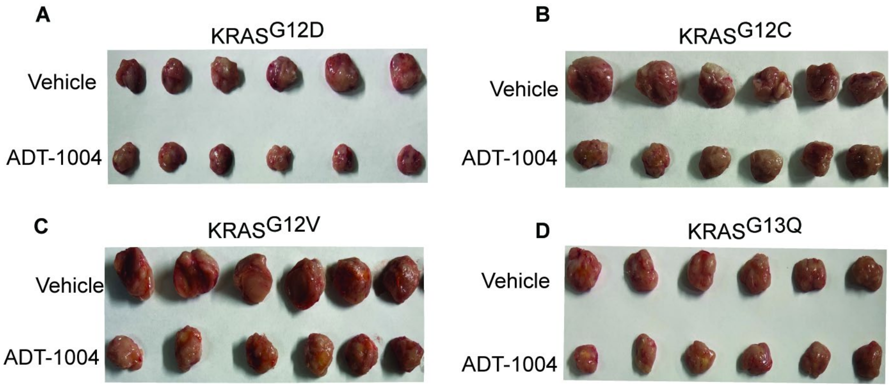


**Supplementary Figure 5: Images of PDX tumors treated with vehicle or ADT-1004 (40 mg/kg).** (**A**). KRAS$^{G12D}$ (**B**). KRAS$^{G12C}$ (**C**). KRAS$^{G12V}$ (**D**). KRAS$^{G13Q}$.

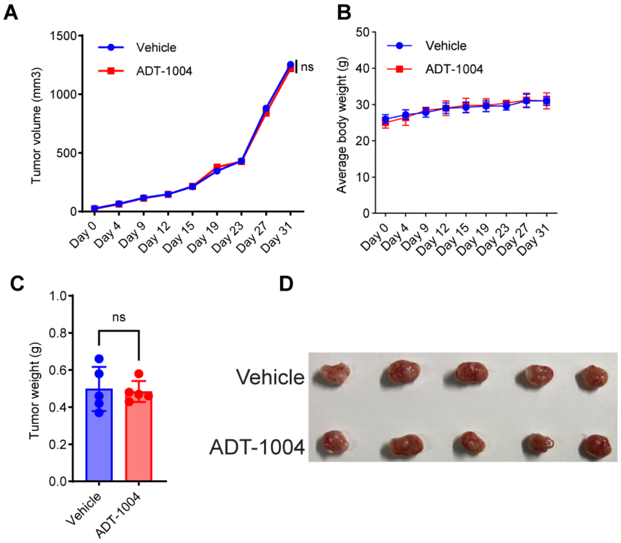


**Supplementary Figure 6: ADT-1004 treatment does not affect tumor growth or molecular markers in the BxPc-3 (RAS$^{WT}$) model.**

(**A**)**.** BxPc-3 cells were subcutaneously injected into the flanks of NSG mice (n=10). Five mice were administered with vehicle or ADT-1004 (40 mg/kg) orally every day for 31 days. The average tumor volume was assessed twice a week and plotted. (**B**). The average body weight of the mice during treatment was measured twice a week. (**C**). Tumor weights were measured from mice at the end of the experiment under the indicated conditions for experiment in (**A**). (**D**) Images of BxPc-3 tumors treated with vehicle or ADT-1004 at the end of the experiment. ns: not significant.

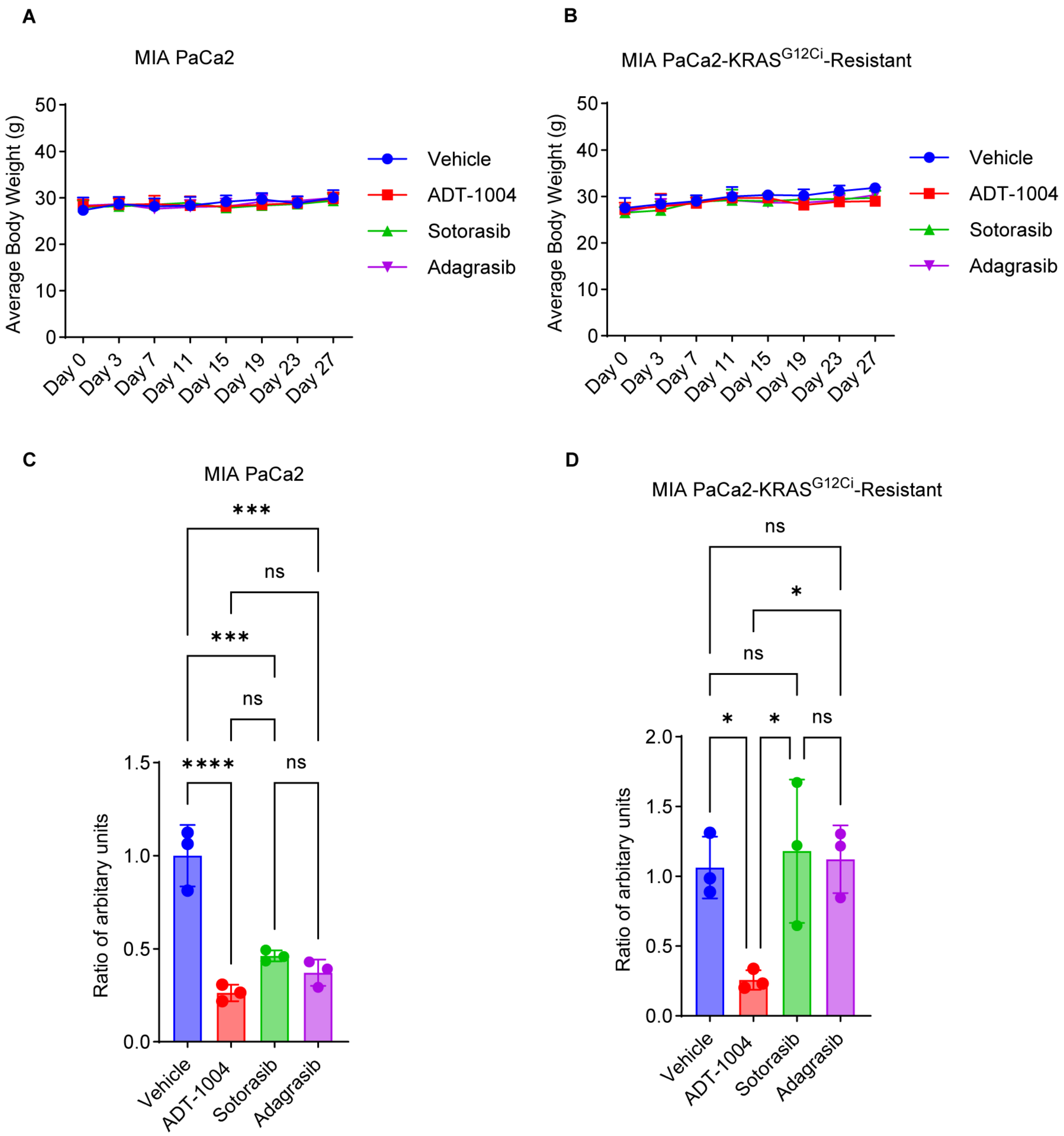


**Supplementary Figure 7: (A).** The average body weights of the mice implanted with MIA PaCa-2 parental cells. **(B).** The average body weights of the mice implanted with MIA-AMG-Res cells. **(C).** Graph depicting the quantification of pERK in Western blot analysis for MIA PaCa-2 parental tumors normalized using β-actin. **(D).** Graph depicting the quantification of pERK by western blot analysis for MIA-AMG-Res tumors normalized using β-actin. ns: not significant, *$p < 0.05$, ***$p < 0.001$, and ****$p < 0.0001$.

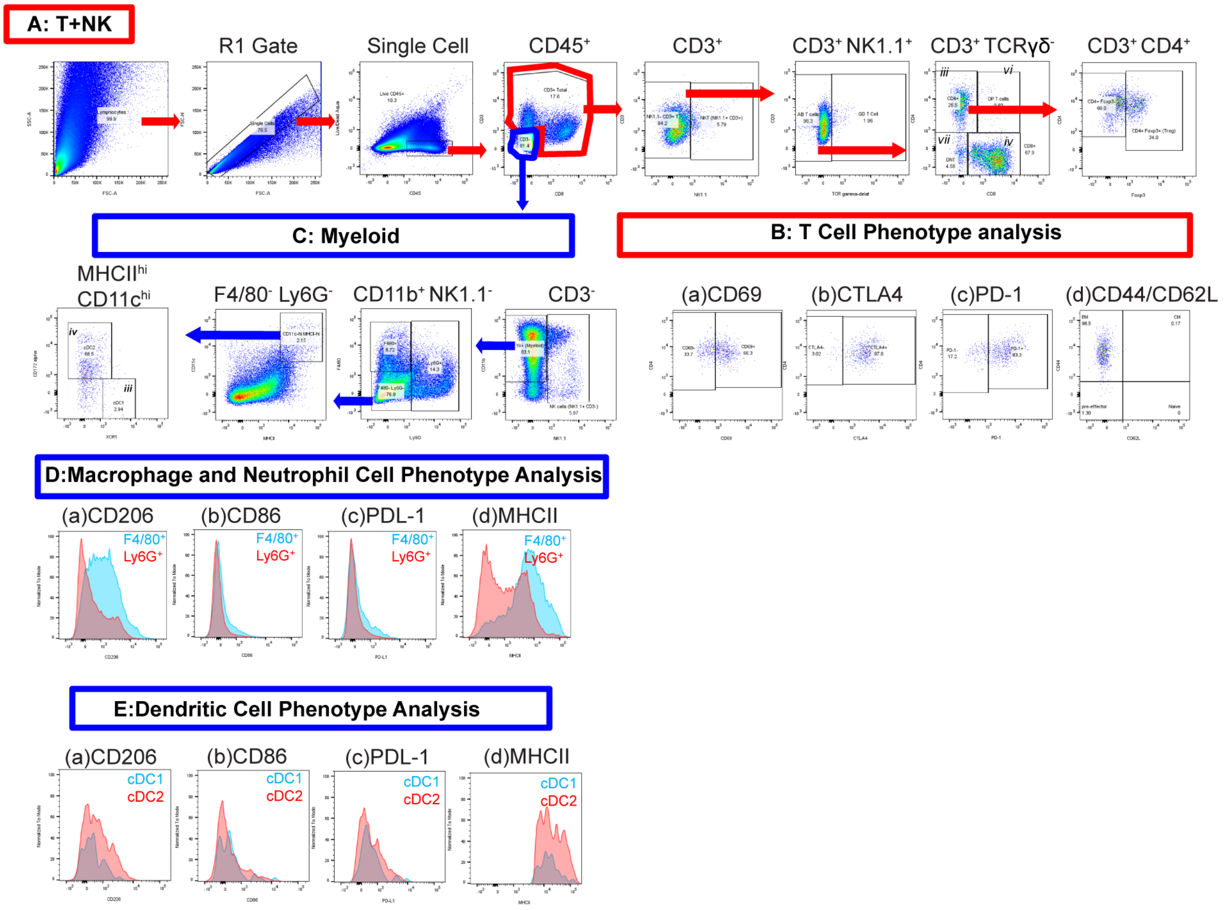


**Supplementary Figure 8. Multiparameter FACS gating strategy for T/NK and myeloid cells.** **(A)** Gating strategy to identify the following T and NK cell subsets: (i) NK T cells ($CD3^+$ $NK1.1^+$), (ii) TCR $\gamma\delta^+$ T cells ($CD3^+$ $NK1.1^-$ $TCR\gamma\delta^+$), (iii) Helper ($CD3^+$ $NK1.1^-$ TCR $\gamma\delta^-$ $CD4^+$) and (iv) Cytotoxic T cells ($CD3^+$ $NK1.1^-$TCR $\gamma\delta^-$ $CD8^+$), (v) Regulatory T cells ($CD3^+$ $NK1.1^-$ TCR $\gamma\delta^-$ $CD4^+$ $Foxp3^+$), (vi) double positive ($CD3^+$ $NK1.1^-$TCR $\gamma\delta^-$$CD4^+$ $CD8^+$), and (vii) double negative ($CD3^+$ $NK1.1^-$TCR $\gamma\delta^-$$CD4^-$$CD8^-$) T cells. **(B)** Phenotypic analysis of T cells was performed for the following markers: (a) CD69, (b) CTLA4, (c) PD-1, (d) CD44/CD62L, (e) CD11b (not shown), (f) TIM-3 (not shown), and (g) LAG-3 (not shown). **(C)** Gating strategy to identify myeloid and dendritic cell subsets, (i) $CD3^-$, (ii) $CD11b^+$ $NK1.1^-$, (iii) $F4/80^-$ $Ly6G^-$, and (iv) $MHCII^{hi}$ $CD11c^{hi}$ **(D-E)** Phenotypic analysis of macrophage, neutrophil, and dendritic cell subsets for the following markers: (a) CD206, (b) CD86, (c) PD-L1, (d) MHCII.

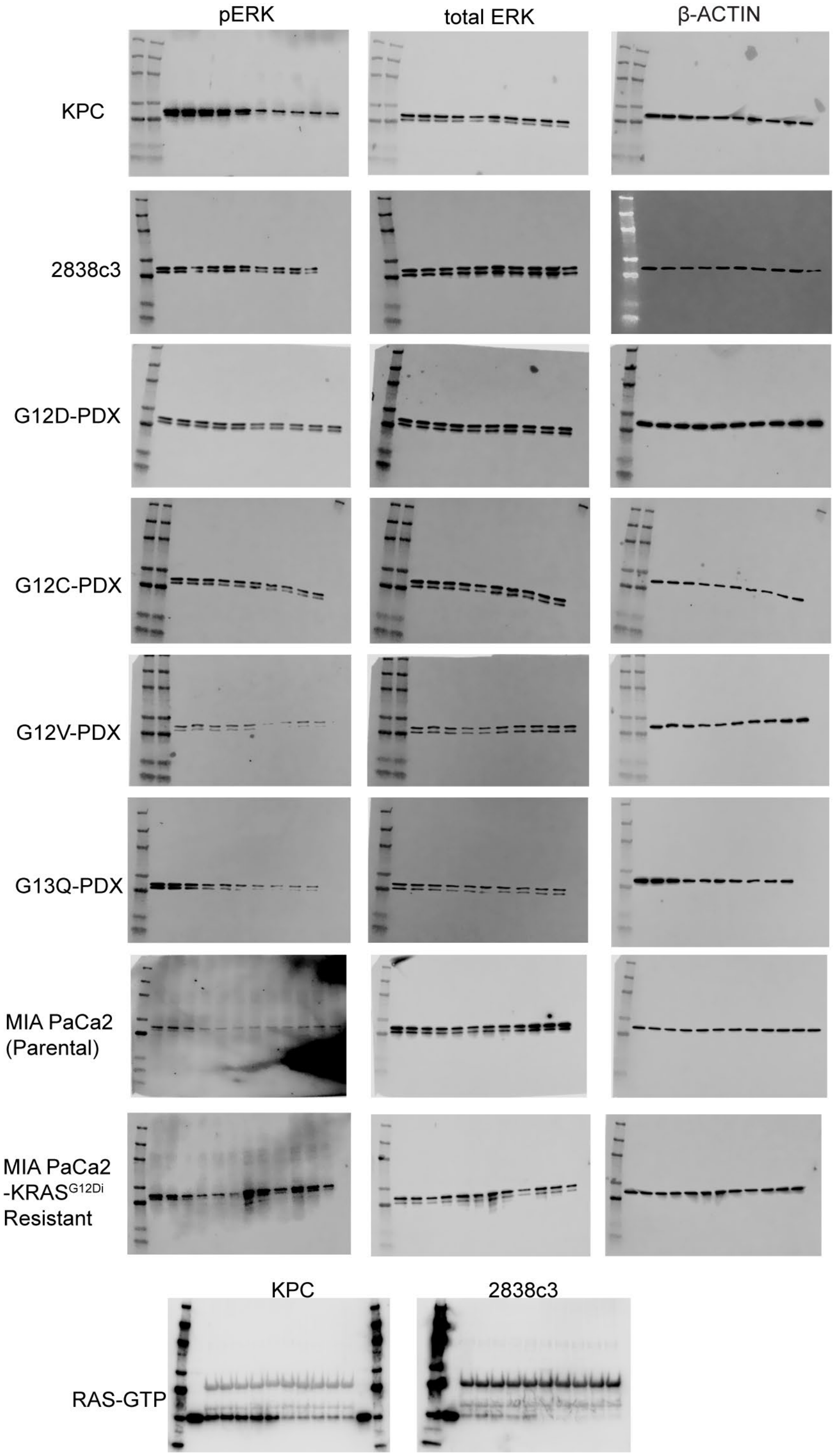


**Supplementary Figure 9:** Raw data of western blots performed in this study.
